# Structure and inhibition mechanisms of *Mycobacterium tuberculosis* essential transporter efflux protein A

**DOI:** 10.1101/2024.09.04.611325

**Authors:** Nitesh Kumar Khandelwal, Meghna Gupta, James E. Gomez, Sulyman Barkho, Ziqiang Guan, Ashley Y. Eng, Tomo Kawate, Sree Ganesh Balasubramani, Andrej Sali, Deborah T. Hung, Robert M. Stroud

## Abstract

A broad chemical genetics screen in *Mycobacterium tuberculosis (Mtb)* to identify inhibitors of established or previously untapped targets for therapeutic development yielded compounds (BRD-8000.3 and BRD-9327) that inhibit the essential efflux pump EfpA. To understand the mechanisms of inhibition by these compounds, we determined the structures of EfpA with inhibitors bound at 2.7 -3.4 Å resolution. Our structures reveal different mechanisms of inhibition for the two inhibitors. BRD-8000.3 binds in a tunnel making contact with the lipid bilayer and extending toward the central cavity to displace the fatty acid chain of a lipid molecule bound in the apo structure, suggesting its blocking of an access route for a natural lipidic substrate, in contrast to its uncompetitive mechanism for the small molecule substrate ethidium bromide which likely enters through an alternative tunnel. Meanwhile, BRD-9327 binds in the outer vestibule without complete blockade of the substrate path to the outside, suggesting its possible inhibition of the dynamical motion necessary for “alternate access” to the two different sides of the membrane, as is characteristic of major facilitator superfamily (MFS) transporters. Both inhibitors may have a role in inhibiting the “alternate access” mechanism that could account for the uncompetitive nature of their efflux of some substrates. Our results explain the basis of the synergy of these inhibitors and their potential for combination in a multi drug strategy for anti-tuberculosis therapy. They also potentially point to a possible function for this essential efflux pump as a lipid transporter. The structures provide a foundation for rational modification of these inhibitors to increase potency.

## Introduction

*Mycobacterium tuberculosis (Mtb)*, the causative agent of human tuberculosis, remains a major global health threat, responsible for 1.3 million deaths in 2022^1^. Mortality rates from tuberculosis are the highest for any bacterial disease^1^. New antibiotics are needed to combat *Mtb* since drug resistance constantly erodes the efficacy of currently used antibiotics. Several first-line antibiotics used against *Mtb*, including isoniazid and ethambutol, affect the cell wall, which provides the first line of defense against the environment within alveolar macrophages. Seeking innovative approaches to target novel vulnerabilities in *Mtb*, we recently developed a broad chemical genetic screening strategy termed PROSPECT (**pr**imary screening **o**f **s**trains to **p**rioritize **e**xpanded **c**hemistry and **t**argets), in which compounds were screened against pools of strains each depleted of a different essential target. Using this approach for generating chemical genetic interaction profiles between small molecules and the set of genetic hypomorphic mutants, we identified new chemical scaffolds that kill *Mtb* by inhibiting novel protein targets including the essential efflux pump A (EfpA)^2^. This strategy enabled our initial identification of BRD-8000 as an inhibitor of EfpA. We performed chemical optimization to achieve potent whole cell activity against wild-type *Mtb* for the analogues BRD-8000.1, BRD-8000.2 and BRD-8000.3, with MIC_90_ of 12.5 µM, 3 µM and 800 nM, respectively (Fig. 1a). Resistance to BRD-8000.3 could be conferred by mutations in EfpA, including substitutions V319F and A415V. BRD-8000.3 is an uncompetitive inhibitor of EfpA efflux of ethidium bromide (EtBr), a demonstrated substrate, although the natural substrate and function of EfpA is not yet identified. Leveraging the same PROSPECT strategy, we subsequently identified a structurally distinct small molecule EfpA inhibitor BRD-9327^3^ (Fig. 1a), based on it sharing a similar chemical-genetic interaction profile with BRD-8000.3. Biochemical evidence revealed that BRD-8000.3 and BRD-9327 work synergistically and display collateral sensitivity with distinct mechanisms of resistance^3^.

**Fig 1.**
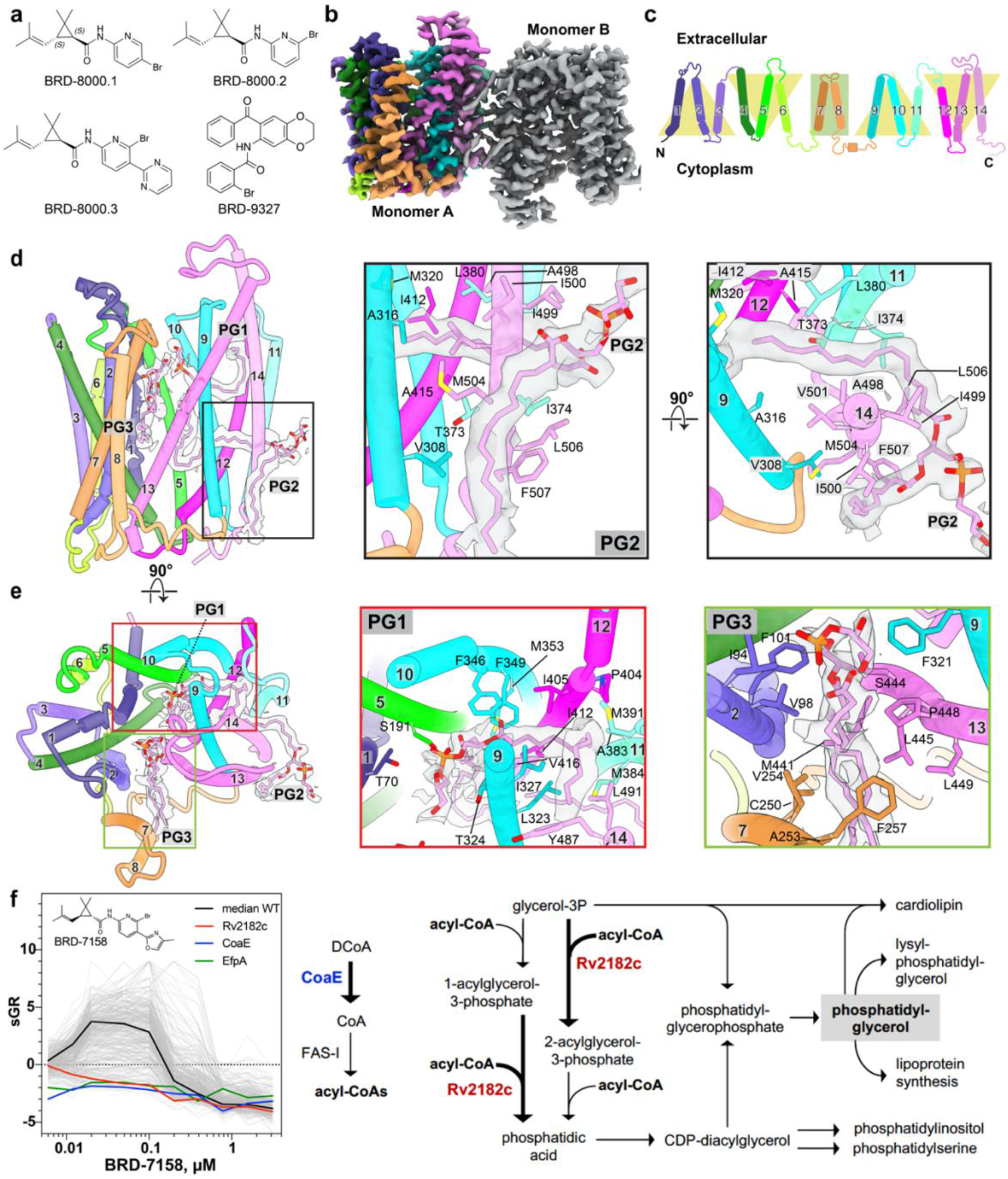
Structure of EfpA. **a,** BRD-8000 series compounds and BRD-9327 identified by PROSPECT assay. **b**, The cryoEM density map for the antiparallel dimer of EfpA^EM^. **c**, Schematic representation of the topology arrangement of EfpA. Three transmembrane helix (™) bundles are related to each other by a twofold pseudosymmetry, with two extra linker helix (™7 and 8). **d**, EfpA monomer A atomic model along with PG molecules (PG1, PG2 and PG3) with corresponding EM density (left panel). PG2 molecule interaction (middle panel) and rotated 90° forward from the position in the middle panel (right panel). The acyl chain of PG2 that is outside of the protein runs parallel to the ™14 hydrophobic residues (F507 ^TM14^, L506 ^TM14^, M504 ^TM14^ and I500 ^TM14^) and reaches downward to the cytoplasmic surface. The head group present between the dimer interface and its second acyl chain enters EfpA via space present between the G377^TM11^, G502^TM14^ residue. This acyl chain is perpendicular to the ™9, ™11, ™12 and ™14 and interact with M320 ^TM9^, A316 ^TM9^, T373 ^TM11^, I374 ^TM11^, L380 ^TM11^,V501 ^TM14^, A498 ^TM14^, I499 ^TM14^, I412 ^TM12^, A415 ^TM12^, V416 ^TM12^ and L419 ^TM12^. **e**, EfpA monomer (left hand panel **D**) rotated 90° forward (left panel). PG1 molecule (middle panel) lies between ™9 (I313^TM9^, A316^TM9^, M320^TM9^, L323^TM9^, I327^TM9^ and Y330^TM9^), ™14 (Y487^TM14^ and L491^TM14^), ™11 (L380^TM11^, M384 ^TM11^ and A383 ^TM11^), ™12 (I405 ^TM12^, P404 ^TM12^,I412 ^TM12^ andV416 ^TM12^), ™10 (F346^TM10^, F349^TM10^and M353^TM10^) ™5 (S191^TM5^) and ™1 (T70^TM1^). PG3 (right panel) lies between ™2 (I94^TM2^, V98^TM2^ and F101^TM2^), ™7 (C250^TM7^, A253^TM7^, V254^TM7^ and F257^TM7^), ™9 (F321^TM9^) and ™13 (M441^TM13^,S444^TM13^, L445^TM13^, P448^TM13^ and L449^TM13^). **f**, Profiles of 361 strains in pooled PROSPECT against BRD-8000 analogue BRD-7158; each strain is shown as a grey line with relevant sensitized strains labeled as in the legend. The x-axis is concentration of BRD-7158 in µM, y-axis is standardized growth rate (sGR) of each strain across 8 doses tested. Right panel shows pathway of PG synthesis highlighting the functions of CoaE and Rv2812c.

EfpA^2^ is an essential transporter in *Mtb* and categorized as a member of the QacA (quaternary ammonium compound A) transporter family^4^. Although EfpA has been suggested to play a role in *Mtb* drug resistance^5^, its essentiality in axenic culture implies that antibiotic efflux is a secondary role, and that the primary function of EfpA is the transport of a yet-uncharacterized biomolecule across the *Mtb* plasma membrane. A better understanding of how the small molecules BRD-8000.3 and BRD-9327 interact with EfpA to prevent transport will support both the potential development of related molecules and the discovery of new classes of inhibitors. Hence, we sought to determine the mechanisms of action of these inhibitors on EfpA by determining structures of EfpA in complex with these compounds.

## Results

### Structure determination of EfpA

*Mtb* EfpA is a 55.58 kDa protein with 14 transmembrane domains (™). We codon optimized EfpA from 3 species of mycobacteria: *Mtb, Mycobacterium marinum* and *Mycobacterium smegmatis* for expression screening in *Escherichia coli* (details in methods section). Most of the structure is contained in the ™ regions with minimal rigid extracellular domains, therefore making it a challenging target for image alignment and structural determination by cryo-EM. To facilitate alignment, we replaced 48 amino acids that preceded the N-terminal ™1 domain and were predicted to be disordered by Alphafold2^6^ by a 12 kDa four-helix soluble domain of BRIL, a variant of apocytochrome b_562_ engineered for stability^7^ (Extended Data Fig. 1a). Fab fragments have been optimized to bind to BRIL providing further prospects for correctly classifying images for three-dimensional reconstruction^8^. Nanobodies that bind to these pre-optimized anti-BRIL Fab fragments have also been optimized and made available, thus providing a common strategy of pre-optimized components that make for efficient structure determination from proteins whose size or structural orientation make them difficult to align^9^. Based on predicted models from Alphafold2 we also introduced a mutation P171R in the loop between ™4 and ™5 of EfpA with the intention that it might act as a site of salt bridge formation with nearby acidic residues (D24, E26, E30, D72 and E71) of the BRIL molecule. The resulting engineered EfpA^EM^ has similar activity to wild-type EfpA as measured by comparison of transport of EtBr, with wildtype EfpA (Extended Data Fig. 1b). The EfpA^EM^ construct was used for protein expression (Extended Data Fig. 1c), and a tripartite complex consisting of purified protein, anti-BRIL Fab and anti-Fab nanobody (Nb) was prepared for cryo-EM^9^ (Extended Data Fig. 1d).

EfpA^EM^ purified as a dimeric assembly, and in single particle analysis of cryo-EM data the dominant arrangement of monomers in the dimer was anti-parallel (Extended Data Fig. 2-3), along with less frequent parallel dimer association (Extended Data Fig. 2). The dimeric assembly may be a result of purification and concentration in detergent lipid micelles. This phenomenon has also been observed in other major facilitator superfamily (MFS) transporters^10,11^. We determined the structures of the antiparallel dimer EfpA^EM^ to 2.7 Å resolution and the parallel dimer to 3.2Å resolution (Extended Data Fig. 2). Complexes were formed with inhibitors BRD-8000.3 and BRD-9327 developed from the chemical genetic screen^2,3^. Their structure was determined in the antiparallel dimer configuration to 3.45 Å and 3.0 Å resolution, respectively (Fig. 2 and Fig.3).

**Fig 2.**
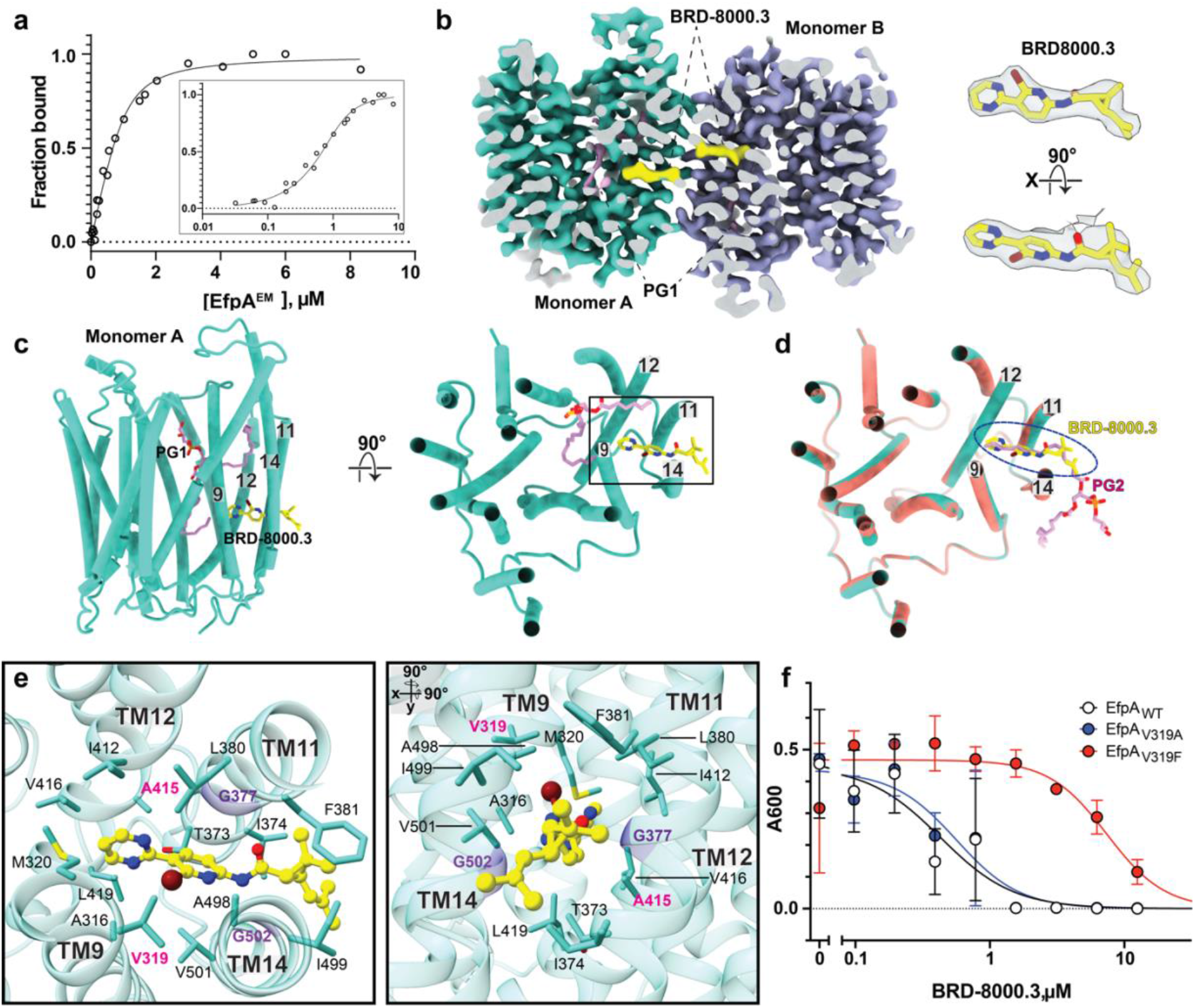
Mode of inhibition of EfpA by BRD-8000.3. **a,** BRD-8000.3 binds to EfpA^EM^ in a ligand-observed NMR assay. Intensity of the inhibitor’s aromatic protons is monitored in a titration series of EfpA^EM^ (x-axis), in which peak heights are converted to fractional occupancy (y-axis). A dissociation constant is determined by fitting the data to a standard bimolecular binding model (see methods). Data represents 3 independent titrations. **b**, Section of cryoEM map of BRD-8000.3 (yellow) bound EfpA^EM^ dimer (left panel). Experimentally determined cryo-EM density of BRD-8000.3 in monomer A (right panel). Map contour level = 0.597 in ChimeraX. **c**, Tertiary structure of the EfpA^EM^ monomer A with BRD-8000.3 (yellow) with lipid shown in pink, side view left and top view from the extracellular side of monomer A. d, Superposition of BRD-8000.3 (yellow) bound EfpA^EM^ monomer (turquoise) onto the apo structure (orange) with PG2 in apo (pink). BRD8000.3 displaces the PG2 molecule present in the apo structure in the BRD8000.3 bound structure. **e**, Tertiary structure for BRD-8000.3 binding pocket showing BRD-8000.3 (yellow) and associated side chains viewed from the extracellular face (left) and side view obtained by rotating left panel at 90° on x and y axis. Entry portal glycines G377^TM11^ and G502^TM14^ are colored violet and side chain atoms colored by atom, nitrogen (blue), oxygen (red), bromide (brown). Residues where mutation led to resistance to BRD-8000.3 are labelled in pink. **f**, Dose-response curves showing growth of *M. bovis* BCG transformed with plasmids overexpressing WT (V319), V319F, and V319A alleles of Mtb EfpA in response to varied concentrations of BRD-8000.3. x-axis is inhibitor concentration and y-axis is culture density after 10 days of growth (OD_600_). Error bars are standard deviation of 3 replicates.

### Overall structure

All four determined structures of EfpA^EM^ are in an outward open conformation with an extracellular gate between TM1, 2, 5 and TM9, 10 open while the intracellular gate was closed by regions of TM4 and TM12, 13 (Fig. 1b). The structure is well ordered for all 14 TM domains and loops (residues 49-518) with exception of 12 disordered amino acids at the C-terminus (Fig. 1b-d). EfpA TM helices organize in a typical MFS transporter fold where two bundles formed by 6 helices (TM1-TM6 and TM9-TM14) are arranged with an internal twofold pseudo symmetry (Fig. 1c). TM7 and TM8 form a linker in the long loop between the two bundles and are juxtaposed next to the TM2 of EfpA (Fig. 1d-e).

In the antiparallel configurations of the dimer seen at 2.7Å resolution, amino acids from TM11 and TM14 of both monomers form the dimer interface. A ‘prime’ after the sequence number indicates amino acids from the second monomer. Y378-Y378’ form pi-pi interactions and I499-I499’ form hydrophobic contacts across the two-fold axis between molecules. These are bolstered by interactions L385-L506’-I374’ above, and L385’-L506-I374 below the two-fold axis on these two helices to form the dimerization interface between molecules (Extended Data Fig. 4a). The structures of the antiparallel dimer are therefore two-fold symmetric assemblies in which the two-fold axis lies in the plane of the bilayer with the same side chains from each molecule displayed in antiparallel arrangement and interdigitated in the interface between molecules above and below the two-fold axis. The parallel dimer uses the same interstitial residues that interdigitate between identical residues in the sequence in a side-by-side manner, resulting in a two-fold symmetric assembly in which the symmetry axis is perpendicular to the membrane plane (Extended Data Fig. 5a-c).

### The structural basis for lipid recognition

Interestingly, we observed extra density in the cryo-EM structures of recombinant EfpA^EM^ protein purified from *E. coli* that is consistent with three, two-chain fatty acid lipids. To identify these lipids, we extracted contents of the micelles and subjected them to lipidomic analysis using normal phase liquid chromatography/mass spectrometry (LC/MS). Lipid peaks were identified as corresponding to phosphatidyl glycerol (PG) and cardiolipin (CDL), with a much less prominent peak corresponding to phosphatidyl ethanolamine (PE), cholesterol hemisuccinate (CHS) and lauryl maltose neopentyl glycol (LMNG) as well as some free fatty acids (Extended Data Fig. 4b). Six well-defined densities line tunnels (numbered 1 to 6) within each monomer and are consistent with aliphatic chains from three diacyl phospholipids. The characteristics of these tunnels were also investigated using the MOLEonline server applied to EfpA after removal of the PG molecules.

The resolution of the head group densities was not clear enough to identify them unambiguously. However, in view of the relatively high abundance of the PG mass spectrometry peak versus PE, we interpreted them as three molecules of PG (PG1, PG2 and PG3) which fit these densities well (Fig. 1d-e and Extended Data Fig. 4c). In theory, the distance between PG1 and PG3 is such that they could be bridged by glycerol to correspond to one, four-chain CDL.

In both the parallel and antiparallel dimers, the density of the two fatty acids chains of PG1 are similar and lie within the protein domain of EfpA^EM^. The phosphatidyl-head group lies in the outer vestibule. One chain lines the wall of the vestibule towards the inner leaflet while the second chain reaches between TM9, TM12, TM13, TM14 toward the outer leaflet. Both fatty acids chains are almost exclusively in contact with hydrophobic side chains from the TMs. The head group is in contact with solvent in the outer vestibule where several polar side chains provide a hydrophilic environment (Fig. 1e).

PG2 densities differed between the parallel and antiparallel dimers. In the antiparallel dimer, one of the fatty acid chains of PG2 penetrates through the side of the protein between TM11 and TM14 away from the external head group towards the central channel. The second fatty acid chain extends from the head group outside the protein downward to the cytoplasmic surface in the region that would be occupied by the inner leaflet. The head group lies outside of the protein in contact with the inner leaflet (Fig. 1d). The head groups of PG2 from each of the two EfpA^EM^ monomers are in contact with each other supporting some degree of hydrophilic contact around the two-fold symmetry axis between them. It is possible that a single cardiolipin (equivalent to two PG2 bridged by glycerol) could fulfill these densities in the micellar complex though this arrangement and the antiparallel dimer itself could not ever be physiological. Two molecules of CHS fill the groove between EfpA^EM^ molecules on the front side (Extended Data Fig. 4c). In contrast, in the parallel dimer, the two fatty acid chains of two individual PG2 molecules of the antiparallel dimer are replaced by a single phospholipid molecule. One acyl chain is inserted in through the sides of each monomer leaving the head group in between monomers (Extended Data Fig. 5a-e). There is extra lipid density (PG4) in the parallel dimer which lies outside the proteins with fatty acid chains reaching toward the cytoplasmic leaflet and replacing the single chains from the two chains of PG2 in the antiparallel dimer.

Finally, in the antiparallel dimer, the PG3 density lies between linking transmembrane domain TM7 and TM8 that connect the two 6-helix bundles, and TM2, TM13 (Fig. 1e and Extended Data Fig. 6a). The head group of PG3 is at the center of the central cavity with acyl chains extending out towards the extracellular leaflet (Fig. 1e). In the parallel dimer, PG3 fatty acyl chains were only weakly defined and were not interpreted.

In order to test the selectivity in the lipid binding sites we performed molecular dynamic (MD) simulations to examine the binding and stability of PG molecules (PG1, PG2 and PG3) in EfpA for a duration of 500 ns using three separate production MD runs. PG1 and PG3 molecules which are completely surrounded by protein exhibit a trajectory RMSD of < 5 Åwhile the PG2 molecule in which one fatty acyl chain is exposed to the surrounding micelle exhibits more motion of that chain with a higher RMSD of < 7 Å(Extended Data Fig. 6b). This is consistent with the tunnels representing stable productive binding sites for lipids within EfpA as seen in our structures.

The presence of PG in the recombinant EfpA protein suggests a pathway for lipid transport that could draw PG or similar lipids from the inner leaflet of the *Mtb* plasma membrane and deliver them to the outer leaflet or the mycobacterial periplasmic space^12^. To gain additional insight, we examined available, expanded PROSPECT data for the BRD-8000 series to define genetic interactions with EfpA inhibition. Specifically, when we examined the chemical genetic interaction profile for a potent analogue, BRD-7158 (MIC_90_ = 0.39 µM; Fig, 1f), against an expanded set of 361 strains, each hypomorphic for a different essential target, we found strong sensitization of additional strains to BRD-7158, beyond EfpA, that are linked to PG synthesis. These highly sensitized strains included Rv2812c (PlsM), an essential acyltransferase, and CoaE, a dephospho-CoA kinase (Fig. 1f). PlsM is an early enzyme in phospholipid synthesis that transfers acyl-CoAs to the *sn-2* position of glycerol-3-phosphate^13^ and *sn*-1-lysophosphatidic acid en route to the synthesis phosphatidic acid (PA) (Fig. 1f), while CoaE executes the final step in coenzyme A synthesis, which is required for the CoA-based activation of the acyl substrates of PlsM^14^. PA is subsequently used for the synthesis of phospholipids including PG, cardiolipin, phosphatidylinositol, and phosphatidylserine^15^. Thus, chemical genetic interactions between EfpA inhibitors and PG synthesis genes support the concept that EfpA may be involved in transporting PG from the inner to outer leaflet of the plasma membrane.

### BRD-8000.3 mechanism of inhibition

We confirmed that BRD-8000.3 binds to EfpA^EM^ using ligand detected proton NMR, with an observed dissociation constant of 179 ± 32 nM (Fig. 2a). The cryo-EM structure of EfpA^EM^ with BRD-8000.3 bound was determined to 3.45 Å resolution (Extended Data Fig. 7). BRD-8000.3 binds in a tunnel (tunnel 2) in EfpA^EM^ that is close to the center of the bilayer (Fig. 2b-c). It displaces PG2 from its position in apo-EfpA with minimal distortion of the surrounding site (Fig. 2d). BRD-8000.3 inserted between TM9, TM11, TM12, and TM14 with its long axis parallel to the membrane plane (Fig. 2c). The BRD-8000.3 binding site shares many of the same residues as the binding site of one fatty acid chain of PG2 and is comprised of hydrophobic amino acid side chains that include A316^TM9^,V319^TM9^, M320^TM9^, T373^TM11^, I374^TM11^,G377^TM11^, L380^TM11^, F381^TM11^ I412^TM12^, A415^TM12^, G413^TM12^, V416^TM12^, L419^TM12^, A498^TM14^, I499^TM14^, V501^TM14^, and G502^TM14^ (Fig. 2e).

Two glycine residues (G377^TM11^ and G502^TM14^) from TM11 and TM14 flank the BRD 8000.3 binding site in tunnel 2 between helices allow access from the bilayer (Fig. 2e center panel). BRD8000.3’s known resistance conferring mutations (V319F or A415V) create steric hinderance to prevent access of BRD-8000.3 to its binding site. To validate this further, we overexpressed WT and mutant EfpA alleles in *Mycobacterium bovis* BCG, an attenuated member of the *Mtb* complex. As expected, overexpression of EfpA mutant V319F resulted in a 13-fold increase in MIC_90_ to BRD-8000.3. However, overexpression of an EfpA allele wherein V319 was changed to a smaller alanine (V319A) had no impact on BRD-8000.3 sensitivity, suggesting that the valine to alanine substitution still allows for inhibitor access and entry to its binding pocket (Fig. 2f).

The structure rationalizes aspects of the structure-activity relationship identified during optimization of the BRD-8000 series as a portent of how it might assist in structure-based optimization of compound affinity. Substitution of the 3-bromo pyridine in BRD8000.1 for 2-bromopyridine in BRD 8000.2 places the bromine substituent into a hydrophobic pocket on just one side of the bromo pyridine ring and produces a ∼4-fold advantage in MIC_90_ ^2^. The further addition of a 1,3-pyrimidine at the 3-position of the pyridine allows extension of the long axis of the molecule further in toward the inside of hydrophobic tunnel 2 and provides a further ∼3-fold advantage in MIC_90_. This direction provides space and hydrophobic structure to support substitution. Resolution of the two trans-stereoisomers of the dimethyl vinyl group in BRD-8000 produced ∼10-fold lower MIC_90_ for the *(S,S)-trans* stereoisomer (BRD 8000.1) than its enantiomer. The active stereoisomer allows favorable hydrophobic interactions with I499, F38, Y378, I374 on the lipid-facing outside of the transporter where this asymmetric environment encodes stereospecificity.

### BRD-9327 mechanism of inhibition

BRD-9327 is a second EfpA inhibitor identified based on similarity of its chemical genetic interaction profile to that of BRD-8000.3^3^ *M. marinum*, a close phylogenetic relative of *Mtb*, mutations that confer resistance to BRD-8000.3 do not confer resistance to BRD-9327 suggesting a different mechanism of inhibition^3^. The structure of BRD-9327-bound EfpA^EM^ at 3.0 Å resolution showed density for BRD-9327 on the inside of the extracellular vestibule with its long axis parallel to transmembrane domains between TM1,TM5,TM9 and TM10 (Fig. 3a-b, Extended Data Fig. 8 and 9a-b). The binding pocket is comprised of T70^TM1^, I73 ^TM1^, V74 ^TM1^, S191 ^TM5^, V192 ^TM5^, L195 ^TM5^, T324 ^TM9^, V325 ^TM9^, I327 ^TM9^, G328 ^TM9^, L329 ^TM9^, V331^TM9^, Q332^TM9^, S338 ^TM10^, A339 ^TM10^, A342 ^TM10^, F346 ^TM10^ and F349 ^TM10^ residues (Fig. 3c).

**Fig 3.**
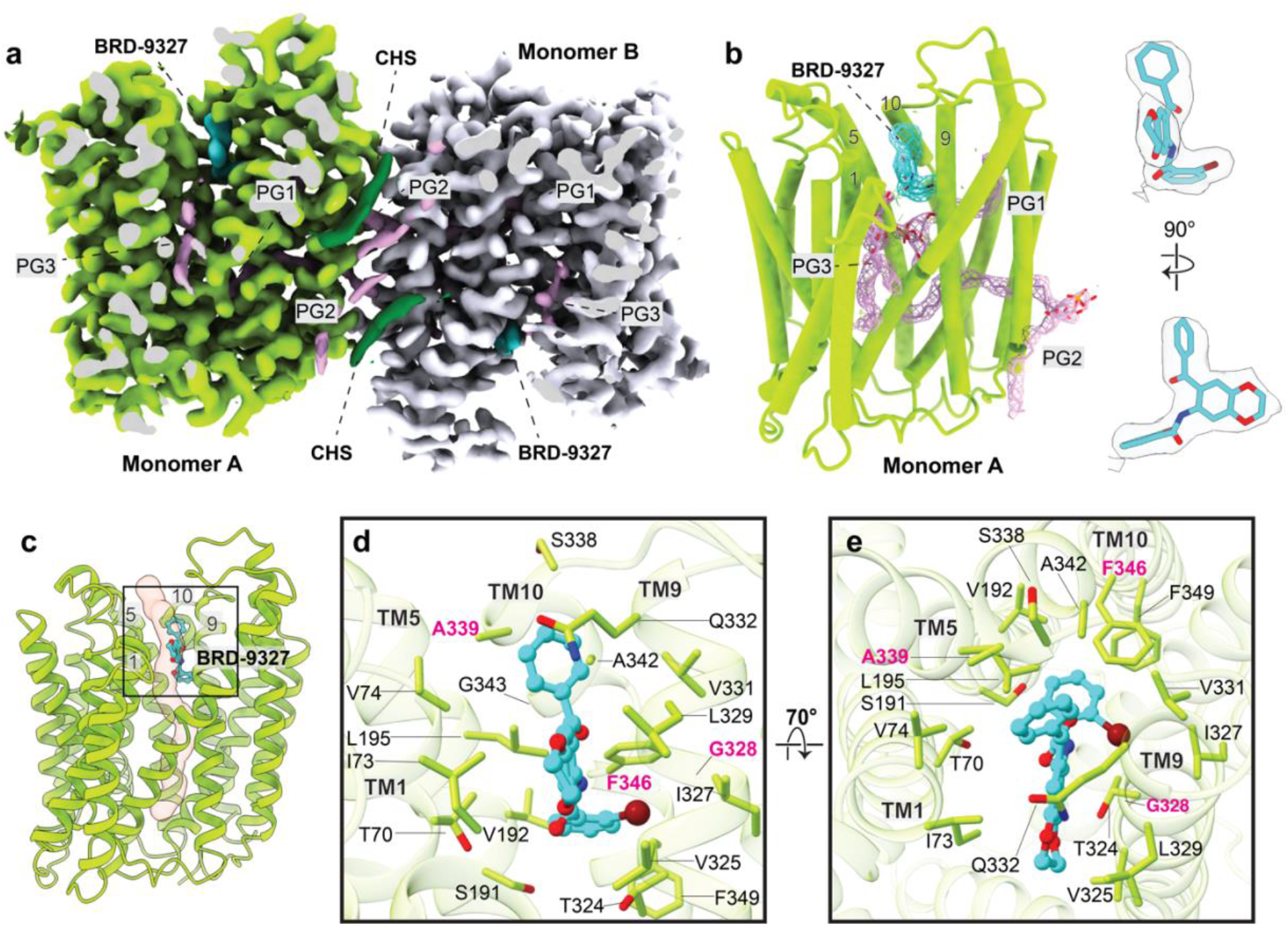
Mode of inhibition of EfpA by BRD-9327. **a**, Sliced cryo-EM map density of BRD-9327 bound EfpAEM. Both monomers of EfpA shown as yellow-green and gray color respectively. Each monomer has one BRD-9327 molecule (aqua color) and three PG molecules (pink color). b, Tertiary structure of EfpAEM monomer A with BRD-9327 (aqua) bound. Density for lipids PG1, PG2 and PG3 are shown in pink (left panel). Cryo-EM density of BRD-9327 (right panel). Map contour level = 0.679 in ChimeraX. c, Structure of BRD-9327 shows it bound in the external vestibule of EfpAEM between TM1,5,9 and 10. d, Model for BRD-9327 binding site in EfpAEM oriented as in c. e, side view rotated by 70° forward from the position in the d. Residues where mutation led to resistance to BRD-9327 are labelled in pink. Color by atom Nitrogen (blue), oxygen (red) and bromide (brown).

Mutations identified to confer resistance to BRD-9327 in *M. marinum* include A339T, G328C, G328D and F346L^3^ (numbering as homologous residue in *Mtb*). These residues are each part of the BRD-9327 binding pocket. Mutations in A339T, G328C and G328D appear to directly interfere with BRD-9327 binding, likely accounting for resistance to the inhibitor. Our MD simulations demonstrate that the phenyl ring of F346 ^TM10^ partially forms π –π interactions with the benzyl bromide of BRD-9327 (Extended Data Fig. 10a-b); a leucine substitution (F346L) which confers resistance would abrogate this interaction. The side chain of Q332^TM9^ above the compound is displaced upward in the presence of BRD-9327 resulting in shift of 2.6Å from the apo state (Extended Data Fig. 9e-f). The head group of PG1 below the molecule and lower in the vestibule is displaced downward away from the compound (Extended Data Fig. 9d-e). The PG1 density is better resolved than in the apo structure, arguably consistent with the closer packing against the compound. The location of BRD-9327 suggests that it will partially block transport of a substrate if it were to pass through the tunnel (Fig. 3c) toward the cytoplasmic side. Also, it will inhibit the dynamical motion required to complete the alternating access transport cycle.

## Discussion

*Mtb* has 30 transporters that are members of the MFS superfamily^12^, only one of which, EfpA, is essential^16^. Interest in EfpA has predominantly centered around its role in drug resistance, with its upregulation in many drug resistant clinical isolates of *Mtb*^17^ and evidence for its ability to efflux a number of first and second line *Mtb* drugs^18^. Meanwhile, the demonstration of its essentiality in the absence of antibiotics both genetically^19-21^ and chemically, with our recent reports of novel EfpA small molecule inhibitors^2,3^, points to a different, critical role for this transporter in bacterial survival and growth. This essential function of EfpA in *Mtb* however has not yet been elucidated, with limited characterization of EfpA outside of the context of drug resistance. EfpA depletion in *Mtb* using CRISPRi has been reported to cause changes in bacterial morphology, possibly suggesting a role in maintaining the cell wall^22^.

Here we sought to determine the structure of EfpA and then to understand the mode of inhibition of two small molecule inhibitors, BRD-8000.3 and BRD-9327 that are not only synergistic, but display collateral sensitivity as resistance to one, confers sensitization to the other^3^. Using a BRIL-based strategy to determine the cryo-EM structure of EfpA, we found that its 14 TM helices are organized in two 6 -helix bundles each (TM1-TM6 and TM9-TM14), arranged with a pseudo twofold symmetry, as is typical for MFS transporters. Structure comparison of the outward open state with other QacA (quaternary ammonium compound A) family members including the *Staphylococcus aureus* QacA transporter shows that EfpA TM arrangement is very similar to QacA (Extended Data Fig. 11). The TM1-6 and TM9-14 of EfpA almost overlap with the QacA transporter while the TM7 TM8 pair are peripheral to the main helical bundles. In QacA^23.^ TM7 TM8 are juxtaposed to space between TM2 and 13 and block the lateral opening while in EfpA TM7 TM8 are arranged opposite, towards TM2, in way that maintains the lateral opening (Extended Data Fig. 11b). This lateral opening is comprised of tunnel 4 and 5 opening into the outer leaflet where a third PG molecule is situated in EfpA (Fig. 4). The lipid transporter MFSD2A also has a lateral opening as in EfpA (Extended Data Fig. 11) where presence of lysophosphatidylcholine has been confirmed in the MFSD2A structure^24^. Presence of this wide lateral opening in EfpA that could accommodate large lipid like molecules, similar to the opening in MFSD2A, suggests that a lipid could be a native substrate for EfpA.

Interestingly, we see lipids occupying the tunnels formed within the recombinant protein isolated from *E. coli*. Despite being purified from *E. coli*, the identification of PG as the dominant co-purified lipid raises the interesting possibility that EfpA may be involved in lipid transport (Fig. 4), possibly for phoshatidylglycerol (PG), a lipid found in all kingdoms of life^25^. In support of this possibility in *Mtb*, we find strong chemical-genetic interactions between genes involved in PG synthesis and EfpA inhibitors, in particular the essential acyltransferase PlsM (Rv2812c) required for the production of phosphatidic acid, a precursor to phospholipids including PG, cardiolipin, phosphatidylinositol and phosphatidylserine^26^. Based on these data, we propose that EfpA could play a role in transport of PG or highly similar lipid.

To propose a model of lipid transport we used Alphafold2 to predict an inward open conformation of EfpA. While all of our cryo-EM structures are in an outward open conformation, we hypothesized the existence of an inward open conformation as is typical of MFS transporters which alternate between inward and outward open conformations as substrates are transported from one face to the other. Alphafold2 predicted another tunnel 1 from the cytoplasmic side (Fig. 4). Thus, substrates such as EtBr, which was the substrate used to evaluate EfpA inhibition^3^ could enter via tunnel 1 while lipid molecules could enter EfpA from the interleaflet region of the plasma membrane via tunnel 1 or tunnel 2 in the inward open conformation. From tunnel 2, the lipid molecule could flop to tunnel 3 and ultimately be transported to the outer leaflet via opening of either tunnel 4 or 6 seen in the outward open structure (Fig. 4). In the gram-negative bacteria *E. coli*, there is a preponderance of PG in the outer leaflet of the membrane in comparison to the inner leaflet^27^. The arrangement of three PG molecules in the structures suggest that EfpA could play a crucial role in maintaining plasma membrane phospholipid asymmetry by transporting the PG from the inner to the outer leaflet (Extended Data Fig. 6a), where it could play a role in membrane organization or fluidity^28^, or serve as a diacylglycerol donor for lipidation of lipoproteins^29,30^. This proposed role would also then be consistent with the morphological changes observed in *Mtb* with knock-down of EfpA^22^.

The inhibitors were uncompetitive with ethidium bromide (EtBr) efflux by EfpA. As they do not obviously block entry at tunnel 1, the presumed entry point of EtBr, or exit from tunnel 6 on the periplasmic side of the membrane, they might achieve this by inhibiting the dynamic transition from inward to outward facing EfpA. This could be the mechanism by which BRD-8000.3 and BRD-9327 inhibit efflux of antitubercular drugs. In contrast, we might expect that BRD-8000.3 has two different ways by which it could inhibit EfpA transport of a natural lipid substrate. As BRD-8000.3 binds in hydrophobic tunnel 2 and displaces one fatty acid chain of the more interior of the PG molecules (PG2) found in the apo structure, it could be competitive with lipid substrates, excluding their entry into tunnel 2 from the lipid bilayer. Additionally, inhibiting the dynamic motions of TM9, TM11, TM12, TM14 could also be key features of the mechanism of BRD-8000.3 inhibition of efflux of a lipid substrate, in a similar manner as that proposed for the small molecule EtBr substrate. In the case of BRD-9327 which binds inside the external vestibule near the external opening of tunnel 6 (Fig. 4). it is uncompetitive with EtBr binding, and does not alter the protein conformation, so does not seem to act as an allosteric inhibitor. Instead, its mode of action could be by blocking either the dynamics of the alternate access mechanism to switch between inner and outer conformations for both natural and unnatural substrates, and/or impeding expulsion of big, natural substrates (larger than EtBr) from the exit portion of the efflux pathway.

**Fig 4.**
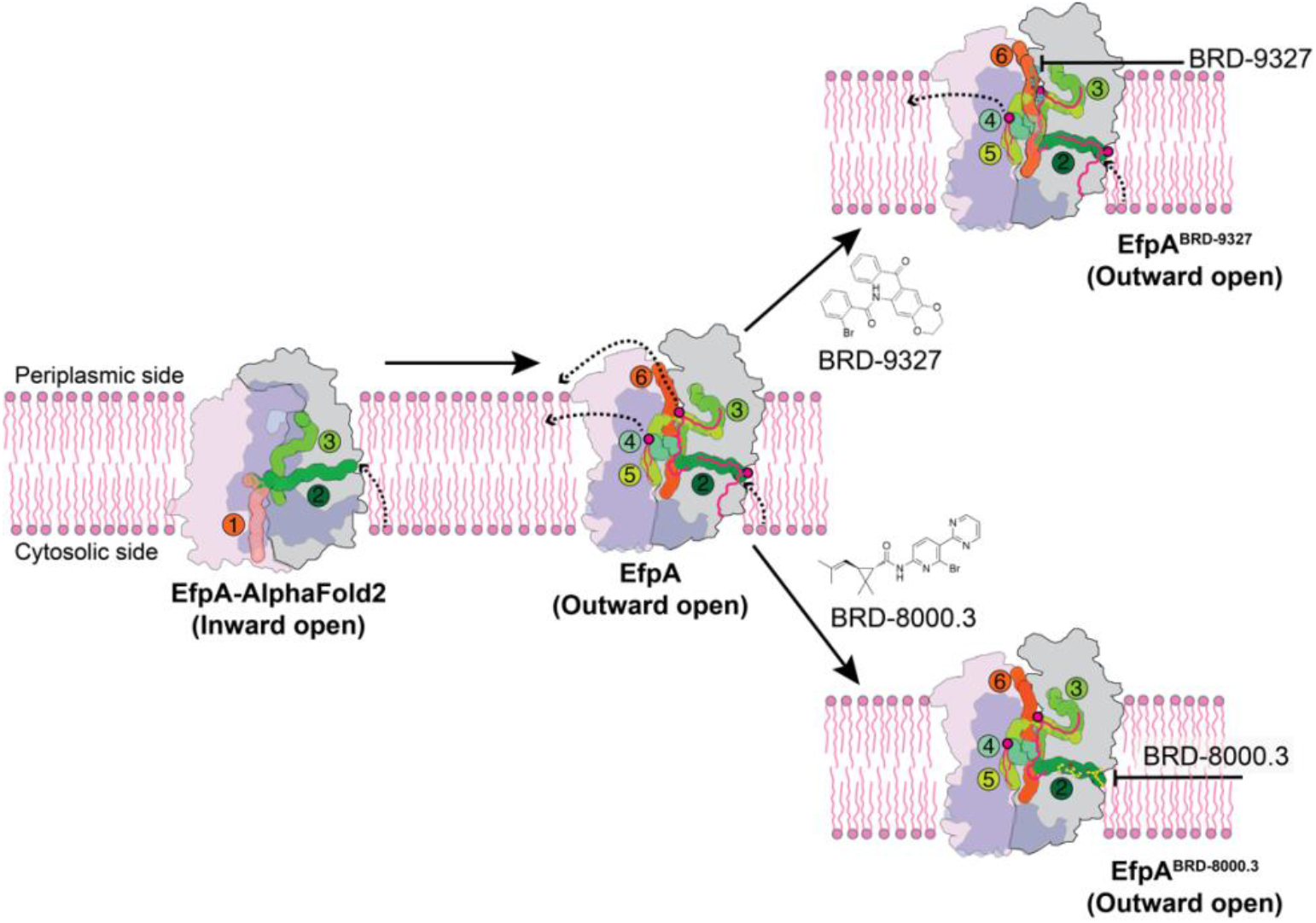
Model of lipid transport by and inhibition of EfpA. The inward facing state was predicted by AlphaFold2. One fatty acid chain of a lipid could bind in tunnel 2, transfer to tunnel 3 for release to the outer leaflet via either tunnel 6 or laterally to the outer leaflet via the 4/5 opening as seen for PG3 in the structure. Tunnels were defined by the MOLEonline server in the inward open model and outward open structure, labeled 1,2,3,4,5,and 6. Schematic showing the binding sites of BRD-8000.3 (bottom) and BRD-9327 (top) to the structure. BRD-8000.3 could be competitive with and block access of the substrate into or from the lipids. BRD-9327 binds in the extracellular vestibule and may interfere with dynamics of the alternate access mechanism, or possibly block export of the true substrate through the periplasmic vestibule in *Mtb*.

In summary the structures we present here uncover different modes of inhibition for inhibitors (BRD-8000.3 and BRD-9327) and support the idea that compounds sharing the same target could be used in combinatorial drug design to treat tuberculosis, potentially with favorable implications for the emergence of resistance^3^. To solidify the hypothesis of possible combinatorial action of these inhibitors, we determined structure of EfpA with both BRD-8000.3 and BRD-9327 exhibiting binding of each at two different locations simultaneously in each monomer transporter (Extended Data Fig. 12). They also provide a crucial platform for the structure-based modification of these compounds to further improve their efficacy. As EfpA is an essential efflux pump in *Mtb*, drugs that inhibit EfpA can inhibit growth of *Mtb* while simultaneously enhancing the activity of companion drugs by limiting their efflux^5,18^, thereby making EfpA an attractive target. Finally, these structures also provide insight into possible functions of EfpA, thereby demonstrating that chemical tools such as these inhibitors can play valuable roles in helping to elucidate the crucial, biological functions of essential targets.

## Acknowledgments

Research was supported by the NIH National Institute of General Medical Sciences (NIGMS GM24485) to RMS. Research to ZG was supported by NIH National Institute of Allergy and Infectious Diseases (NIAID R01AI178692). Research to DTH was supported by the Bill and Melinda Gates Foundation. We thank Anthony Kossiakoff for provision of the anti-BRIL Fab, and nanobody against it. We thank Abhijit A. Sardesai for gift of *E. coli* JD838 strain. We thank Sun Kyung Kim for help in expression cloning. We thank D. P. Bulkley, G. Gilbert and M. Harrington for support with cryo-EM data collected at the UCSF cryo-EM facility, which is supported by NIH grants S10OD020054, S10OD021741 and S10OD026881. We thank Janet Finer-Moore for critical reading of the manuscript.

## Author contributions

N.K.K., D.T.H., and R.M.S. conceived the project. N.K.K. and M.G. prepared the protein. M.G. and N.K.K. prepared EM grids, collected data and processed cryo-EM data. N.K.K. and M.G. performed model building and refinement. Z.G. determined the lipids by mass spectrometry. S.B. performed the NMR study. J.E.G. and A.Y.E. performed the PROSPECT analysis and MIC testing in BCG. T.K. performed chemical synthesis. S.G.B. and A.S. performed the MD-simulation analysis. N.K.K., M.G., J.E.G., D.T.H., and R.M.S. wrote the manuscript with contributions from all other authors.

## Data and materials availability

Data availability: The atomic coordinates for four structures of EfpA apo antiparallel dimer, parallel dimer, BRD-9327 bound and BRD-8000.3 bound have been deposited in the Protein Data Bank with the accession codes PDB ID 9BII, 9BL7, 9BIQ and 9BIN respectively. The corresponding maps have been deposited in the Electron Microscopy Data Bank with the accession codes EMD-44591, EMD-44651, EMD-44598 and EMD-44594.

## Materials and Methods

### Protein expression and purification

The *Mycobacterium tuberculosis* EfpA (Efflux protein A) protein (Uniprot: P9WJY4) encoding gene along with N-terminal Flag (DYKDDDK) epitope tag and C-terminal 10X histidine tagged fusion protein was codon-optimized and cloned into the pET-28a(+) bacterial expression vector (Twist bioscience). The BRIL-tag construct was prepared from it by replacing the first 48 amino acids of EfpA that are predicted to be disordered by Alphafold2 and lie in the cytoplasmic space with BRIL (a variant of apocytochrome b562)^7,31,32^ to serve as a fiducial marker. The final EfpA-EM construct was derived from BRIL-EfpA by mutating proline at position 171 to arginine (GenScript) to provide possible interactions that might help order the BRIL domain.

The resulting construct pET-28a-BRIL-P171R_EfpA was transformed into *Escherichia coli* BL21(DE3) strain (NEB) and a primary culture was set from the colony obtained after transformation. From primary culture secondary culture set in 4 flask of TB media 1 L each with 50 μg/mL kanamycin and grow at 37 °C, 200 rpm until O.D. 600nm. reach between 0.6-0.8. Induction was performed by adding 0.5 mM of IPTG and grown for another 4 hours 37 °C followed by harvesting with centrifugation and stored at −80 °C.

For protein purification, cells were resuspended in ice cold lysis buffer (50 mM Tris-Cl pH 8.0, 500 mM NaCl, DNase I and cOmplete, EDTA-free protease inhibitor cocktail tablets (Roche)) and lysed using sonication pulse (1s on and 1s off) on ice for 3 minutes in 5 cycles. The lysate was spun at 112,967 × g for 2 hours ultracentrifugation to separate the crude membrane and membrane pellet and stored at −80 °C. Detergent extraction of protein was performed by resuspension of membrane (15 mL buffer/1 g membrane) in ice cold resuspension buffer (50 mM Tris-Cl pH 7.5, 300 mM NaCl, 0.5% 2,2-didecylpropane-1,3-bis-β-D-maltopyranoside (LMNG)/0.05% cholesteryl hemisuccinate (CHS) and protease inhibitor) at 4 °C overnight. Insoluble material was separated by centrifugation at 34,155 × g for 30 min at 4 °C and supernatant was filtered through a 0.4 µM filter.

Filtered soluble protein supplemented with 30 mM Imidazole pH 7.5 and loading was performed at a flow rate of 1.5 mL/min onto a 5 mL HisTrap HP column (Cytiva). The column was washed extensively first with Buffer A (50 mM Tris-Cl pH 7.5, 300 mM NaCl, 0.01% LMNG/0.001% CHS) followed by buffer A supplemented with a gradient of imidazole, concentration (30mM, 50mM, 80mM, 120mM). Protein was eluted in 20 mL elution buffer (50 mM Tris-Cl pH 7.5, 300 mM NaCl, 300 mM imidazole, 0.01% LMNG/0.001% CHS. Protein was concentrated in a 50-kDa filter (Amicon) and purified by injecting in Superdex 200 Increase 10/300 GL column (Cytiva) equilibrated with buffer A. Fractions containing EfpA protein were concentrated, flash frozen in liquid nitrogen and stored at −80 °C.

### Anti-BRIL Fab (BAG2 Fab) purification

The BAG2 Fab expression construct in pRH2.2 vector was a gift from Kossiakoff^8^. This vector was transformed into competent C43(DE3*) E. coli* cells (Sigma-Aldrich) and grown overnight in 10 mL 2xYT supplemented with 100 μg/mL ampicillin shaking at 200 rpm and 37 °C. 10 mL overnight culture was used to start secondary culture in 1L of sterile TB autoinduction media (Terrific Broth supplemented with 0.4% glycerol, 0.01% glucose, 0.02% lactose and 1.25 mM MgSO4 and 100 μg/mL ampicillin) in a 2.8 L baffled flasks at 37 °C for about 6 hours and then decrease the temp to 30 °C and leave it overnight at 225 rpm. Next day cells were harvested by centrifugation and stored in the −80 °C. Purification was performed by lysis of cells using sonication in 50 mM Tris-Cl pH 7.5, 300 mM NaCl supplemented with protease inhibitors, DNase I, and the soluble fraction was heated in a 60 °C water bath for 30 min. High-speed centrifugation was performed to remove aggregates and the supernatant affinity purified using protein L purification (Cytiva) with the manufacturer’s protocol. The eluted Fab was concentrated with a 10 kDa MWCO spin and further purified with size-exclusion chromatography, using a Superdex S75 Increase 10/300 column (Cytiva) equilibrated in 50 mM Tris-Cl pH 7.5, 300 mM NaCl and purified sample was flash freeze in liquid N2 and stored in −80 °C.

### Expression and purification of anti-Fab elbow Nb

DNA encoding Nb^33^ was cloned into a pET-26b expression vector with N-terminal 6x His and Nb sequence. The expression vector was transformed into BL21(DE3) cells (NEB), and primary culture started in 50 mL 2xYT. A secondary culture was started in 6 flasks with 1 L 2xYT each with 50 μg/mL kanamycin and grown at 37 °C until mid-log (0.6-0.8) phase and induced with 1 mM IPTG. Expression was carried out overnight at 25°C. Culture was pelleted down and lysed by sonication in 100 mL lysis buffer (50 mM Tris, 500 mM NaCl, protease inhibitors and DNase I). The lysate was spin down at 20,000g for 45 min and supernatant filtered using a 0.4-micron filter. The sample was supplemented with 5mM imidazole and loaded onto 5 mL HisTrap HP column (Cytiva). The protein was purified from the Ni column by the protocol used for the EfpA purification. Purified protein sample were concentrated up to 5 mL using a 3K centrifugal filter and further purified using a Superdex S75 Increase 10/300 column (Cytiva), equilibrated in 50 mM Tris-Cl pH 7.5, 300 mM NaCl and stored in −80 °C.

### Cryo-EM sample preparation and data collection

For cryo-EM sample preparation, the purified EfpA-EM protein, anti-BRIL Fab, and anti-Fab elbow Nb were mixed in the ratio of 1:1.5:2 and incubated for 1 h. on ice. After incubation the complex were separated using a Superose 6 Increase 10/300 GL column equilibrated with 50 mM Tris-Cl pH 7.5, 300 mM NaCl, and 0.02% GDN. The peak fractions corresponding to the ternary complex of EfpA-EM protein with Fab and Nb were collected (Extended Data Fig.1) and concentrate before use for cryo-EM grid preparation.

For EfpA apo structure 4 μL of protein complex sample at concentration of 16.16 mg/mL was applied to cryo-EM grids Quantifoil AU R1.2/1.3, 400 mesh (Electron Microscopy Sciences) using Mark IV Vitrobot (FEI) at 8 °C, 100% humidity and blotted for 3.5 s followed by plunge-freezing in liquid ethane.

For BRD-8000.3 sample the protein was incubated with BRD-8000.3 100 μM final concentration on ice for 15 minutes and 4 μL of sample of final 5.91 mg /mL protein with 100 μM BRD-8000.3 was applied on cryo-EM grids Quantifoil AU R1.2/1.3, 400 mesh (Electron Microscopy Sciences) before being plunged into liquid ethane equilibrated to -185 ºC.

A total of 16,938 movies were recorded for EfpA apo samples on a Titan Krios at 300 kV equipped with a K3 Summit detector (Gatan) using SerialEM v4.1 beta software at UCSF. Data were collected in non-super resolution mode at 105,000X magnification (pixel size of 0.835 Å/pix) with a defocus range of -1 to -2 µm. Each movie contains 80 frames with a total electron dose of ∼45 electrons /Å^2^.

A total of 10,235 movies were recorded EfpA-BRD-8000.3 samples on a Titan Krios at 300 kV equipped with a K3 Summit detector (Gatan) using SerialEM v4.1 beta software at UCSF. Data were collected in super resolution mode at 105,000X magnification (pixel size of 0.4175 Å/pix) with a defocus range of -1 to -2 µm. Each movie contains 80 frames with a total electron dose of ∼45 electrons /Å^2^.

For BRD-9327 bound sample protein was incubated with 200 μM final concentration of BRD-9327 on ice for 15 minutes and 4 μL of sample of final 8.9 mg /mL protein with 200 mM BRD-9327 was applied on cryo-EM grids Quantifoil AU R1.2/1.3, 400 mesh (Electron Microscopy Sciences) before being plunged into liquid ethane equilibrated to -185 ºC. A total of 12,748 movies were recorded on a Titan Krios at 300 kV equipped with a K3 Summit detector (Gatan) using SerialEM v4.1 beta software at UCSF. Data were collected in super resolution mode at 105,000X magnification with physical pixel size of 0.417 Å/pix and defocus range of -1 to -2 µm. Each movie contains 80 frames with a total electron dose of ∼43 electrons /Å^2^.

### Cryo-EM data processing

Raw movies were motion-corrected using UCSF MotionCor2v1.4.1^34^. For BRD-8000.3 and BRD-9327 super resolution mode motion-corrected data to Fourier bin images 2 × 2 to the counting pixel size 0.835 Å/pix and 0.834 Å/pix respectively. Dose-weighted micrographs were imported into cryoSPARC^35^ v4.2.1 and defocus values estimated using patch CTF estimation (multi-GPU). Particle picking, extraction, classification and refinement were performed in cryoSPARC v4.2.1 as depicted in workflows shown in extended data (Extended Data fig. 2, 7 and 8).

### Model building and refinement

A model of EfpA from AlphaFold2 was used as starting model and rigid body and manual fitting was performed in COOT^36^ followed by real-space refinement jobs in by Phenix ^37,38^ against the final map. BRD-8000.3, BRD-9327, PG and CHS restraints were generated using phenix eLBOW. The final model statics are reported in Extended Data Table 1. Figures were prepared using UCSF ChimeraX^39,40^.

### MIC assays

The efpA coding region from *M. tuberculosis* H37Rv was PCR amplified using primers efpA_pUV_clone_F1 (GTTAATTAAGAAGGAGATATACATATGACGGCTCTCAACGACACAGAG) and efpA-pUV-R1 (GAATATTACAGCTCGCCGGCGTCGAT) and cloned into the PCR4 TOPO vector using the Invitrogen TOPO TA cloning kit (REF 45-0071). V319F and V319A mutations were introduced using Agilent QuikChange II Site-Directed Mutagenesis Kit (Cat #200524). WT and mutant *efpA* regions were excised from the TOPO vector using SspI and PacI and cloned into the mycobacterial shuttle vector pUV15tetORm digested with PacI and EcoRV. Resulting constructs were transformed in *Mycobacterium bovis* BCG (Pasteur).

BCG was grown in Middlebrook 7H9 medium (Difco) supplemented with 10% OADC (BBL), 0.2% glycerol, 0.05% tyloxapol, and 50 µg/ml hygromycin B throughout the experiment. Transformants were grown to mid log phase (A600mn 0.2-0.3) and EfpA overexpression was induced 1d prior to MIC assay with 100 ng/ml anhydrotetracycline (AHT). Following overnight induction, cultures were back diluted to A600nm 0.0025 prior to transfer into 96 well assay plates (100 µl/well) containing 1 µl/well of BRD-8000.3 serially diluted in DMSO. Plates contained 3 replicates at each dilution and were incubated at 37°C for 10 days.

### EtBr efflux assay

Wild type EfpA and EfpA^EM^ genes were cloned into a pBAD vector and transformed into JD838 cells, a strain derived from *E*. *coli* K-12 in which three drug transporters are deleted (genotype: Δ*mdfA*Δ*acrB*Δ*ydhE::kan*). The JD838 strain has been previously used for EtBr transport assay with different MFS transporters^41^. The bacterial cultures were grown at 37 ºC in luria bertani (LB) media to an optical density (OD_600_) of 0.4 before induction of EfpA expression with 0.05% arabinose. After 2 hours, the cells were harvested, washed with PBS, and the pellet resuspended in EtBr-free PBS by adjusting the OD_600_ to 1. The cells were then loaded with 20 μM Ethidium Bromide (EtBr) in the presence of 0.5 μM of carbonyl cyanide m-chlorophenyl hydrazone (CCCP) and incubated in the dark at 37 ºC for 45 minutes. After incubation, the cells were washed three times and resuspended in PBS again by adjusting the OD_600_ to 0.8. EtBr efflux was initiated by adding 50 μL of cells into a 96 well clear bottom plate containing 50 μL 0.8 % glucose in PBS. EtBr fluorescence was then monitored continuously for 45 minutes using a SpectraMax M5 plate reader (Molecular Devices) preset to an excitation wavelength of λ = 530 nm and emission wavelength λ = 585 nm.

### Extraction of lipids co-purified with MFS transporter protein

Lipids co-purified with the protein sample were extracted using the method of Bligh and Dyer as previously described^42^. Briefly, the protein solution (containing ∼1mg protein) was transferred into a glass tube, and phosphate-buffered saline (PBS) solution was added to a final volume of 1.6 mL. Then, 2 mL of chloroform and 4 mL of methanol were added to make the single-phase Bligh-Dyer mixture, which consists of chloroform/methanol/PBS (1:2:0.8, v/v/v). This solution was subjected to sonic irradiation in a bath apparatus for 5 min. This single-phase extraction mixture was then centrifuged at 500x g for 10 min in a clinical centrifuge to pellet the protein precipitate. The supernatant was then transferred to a fresh glass tube where 2 mL of chloroform and 2 mL of PBS were added to generate the two-phase Bligh-Dyer mixture, which consists of chloroform/methanol/PBS (2:2:1.8, v/v/v). After mixing and centrifugation (500x g) for 10 min, the upper phase was removed, and the lower phase was dried under a stream of nitrogen. The dried lipid extract was stored at -20 °C until LC/MS analysis.

### Lipid Identification by LC/MS/MS

Normal phase LC was performed on an Agilent 1200 Quaternary LC system equipped with an Ascentis Silica HPLC column, 5 μm, 25 cm x 2.1 mm (Sigma-Aldrich, St. Louis, MO) as previously described^43^. Mobile phase A consisted of chloroform/methanol/aqueous ammonium hydroxide (800:195:5, v/v/v); mobile phase B consisted of chloroform/methanol/water/aqueous ammonium hydroxide (600:340:50:5, v/v/v); mobile phase C consisted of chloroform/methanol/water/aqueous ammonium hydroxide (450:450:95:5, v/v/v). The elution program consisted of the following: 100% mobile phase A was held isocratically for 2 min and then linearly increased to 100% mobile phase B over 14 min and held at 100% B for 11 min. The LC gradient was then changed to 100% mobile phase C over 3 min and held at 100% C for 3 min, and finally returned to 100% A over 0.5 min and held at 100% A for 5 min. The LC eluent (with a total flow rate of 300 μm/min) was introduced into the ESI source of a high resolution TripleTOF5600 mass spectrometer (Sciex, Framingham, MA). Instrumental settings for negative ion ESI and MS/MS analysis of lipid species were as follows: IS= -4500 V; CUR= 20 psi; GSI= 20 psi; DP= -55 V; and FP= -150 V. The MS/MS analysis used nitrogen as the collision gas. Data analysis was performed using Analyst TF1.5 software (Sciex, Framingham, MA).

### Molecular dynamics simulations

All atom molecular dynamics (MD) simulations were performed with the GROMACS simulation engine^44^ using the CHARMM36 force field^45^ parameters. The cryoEM structure was used as the starting point for the MD simulations. CHARMM-GUI^46^ was used to prepare the system by embedding the protein in a lipid bilayer which was modeled using POPC molecules. The initial membrane coordinates were designated by the PPM server through the Charmm-GUI interface. The system was further solvated using the TIP3P water model with the inclusion of 150 mM of Na^+^ and Cl^-^ ions. The solvated system was minimized for 5000 steps using the steepest descent method. Furthermore, the system was equilibrated with default parameters supplied by Charmm-GUI, starting with weak restraints, and successively reducing the strength of the restraints in 5 steps for a total time of 50 ns. Additionally, a 100 ns equilibration simulation was run without any restraints. Following this, production simulations were executed for 500 ns within the NPT ensemble, employing a Parrinello– Rahman barostat set to 1 atm and a V-rescale thermostat maintaining a temperature of 303.15° K. The SHAKE algorithm was applied to enforce constraints on all bond lengths involving hydrogens and the hydrogen mass repartitioning scheme was utilized for the production runs, enabling a time step of 4 fs.

### BRD-8000.3 binding assay

1H-NMR experiments were recorded on a Bruker 600MHz spectrometer equipped with a cryogenic QCIF probe and an automatic sample handling system. Experiments were conducted either at 280K or 298K and processed using the spectrometer’s TopSpin 3.6 software. Intensities of the most prominent aromatic proton peak ∼8.9ppm were extracted in a titration series with and without protein and normalized to reflect free and bound BRD-8000.3. Samples typically contained 30 µM BRD-8000.3 in NMR buffer (50mM Tris-d7 300mM NaCl, 0.01%LMNG and .001%CHS at pH7). The data were fit to a standard quadratic binding model to determine the apparent affinity^47^.

### PROSPECT of BRD-7158

PROSPECT was executed as described^2^. BRD-7158 was among 21 compounds with known targets profiled in 10-point dose response using an expanded 361-strain PROSPECT pool (354 hypomorphic strains and 7 barcoded H37Rv strains). After initial calculation of strain-specific log2-fold changes as described previously ^2^ train-by-strain effects were converted to growth rate (GR, 1 = no effect, 0 = full effect) based on comparison to strain behavior in DMSO and rifampin (100nM) respectively. GR for each strain was then standardized (z-scored) across all test treatments in the screening data, generating a standardized growth rate (sgr) metric that provides confidence in the scale of the measured effect.

