## Supplementary Figures for "Structure and inhibition mechanisms of *Mycobacterium tuberculosis* essential transporter efflux protein A": Extended_data_Khandelwal_Gupta_etal.pdf

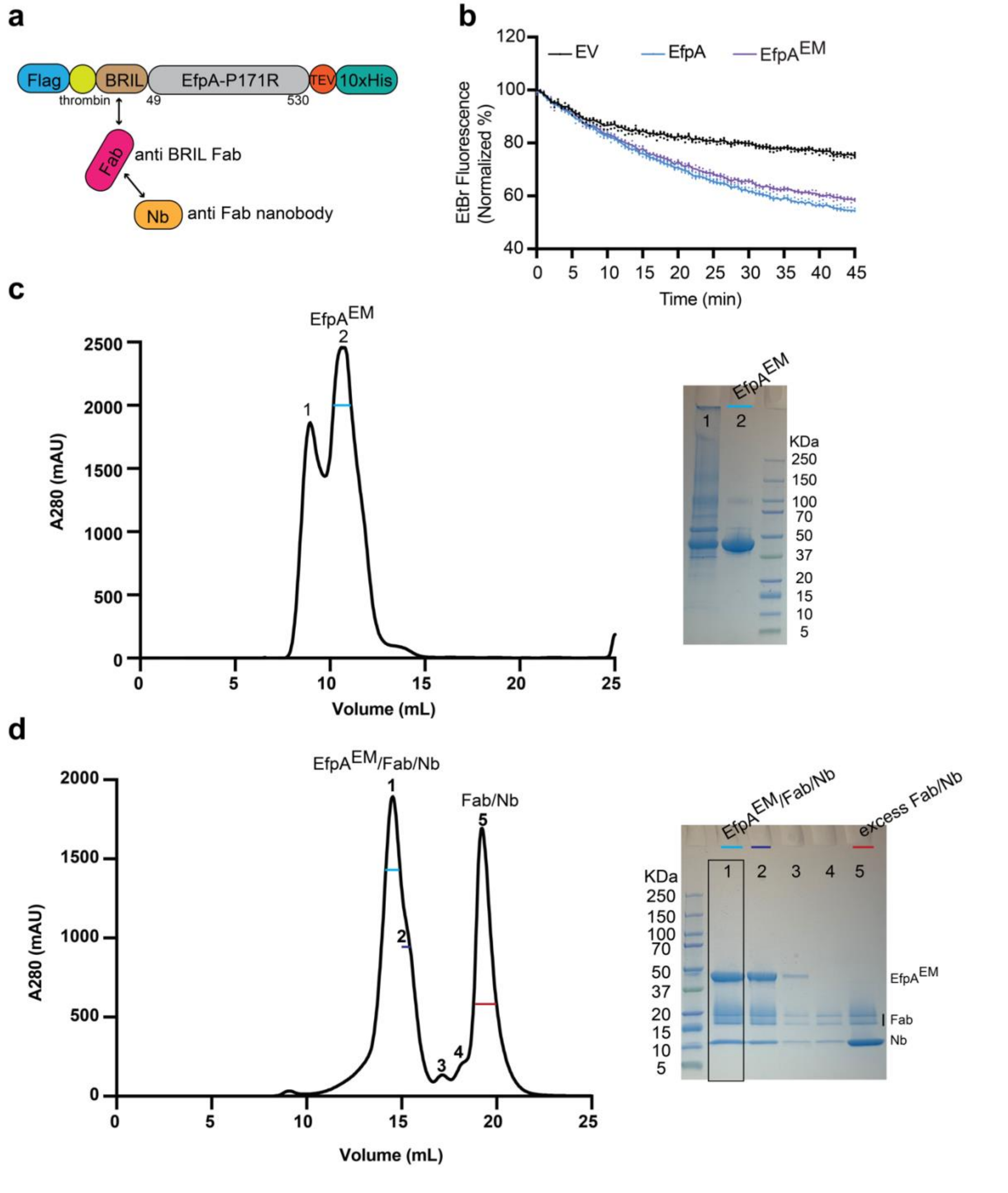


**Extended Data Fig. 1| EfpA$^{EM}$ construct design and purification. a,** Construct design and complex formation strategy for EfpAEM. **b,** Whole cell-based transport of EtBr by EfpA. *E. coli* JD838 strain (Δ*mdfA*Δ*acrB*Δ*ydhE::kan*) cells expressing wildtype EfpA (light blue), EfpA$^{EM}$ (purple) and empty vector (black) were pre-loaded with 20 μM EtBr. After stimulation with glucose, ethidium fluorescence intensity decay at 585 nm was monitored continuously using a plate reader for 45 minutes. All points for experiments (n = 4) are shown in the graph with average showing as solid line. **c,** Sizing profile of $EfpA^{EM}$ and SDS gel. Numbers on the image correspond to labels on the peaks in the sizing profile. **d,** Sizing profile of purified $EfpA^{EM}$ /Fab/Nb complex and SDS gel. Peak 1 was used for cryoEM grid preparation.

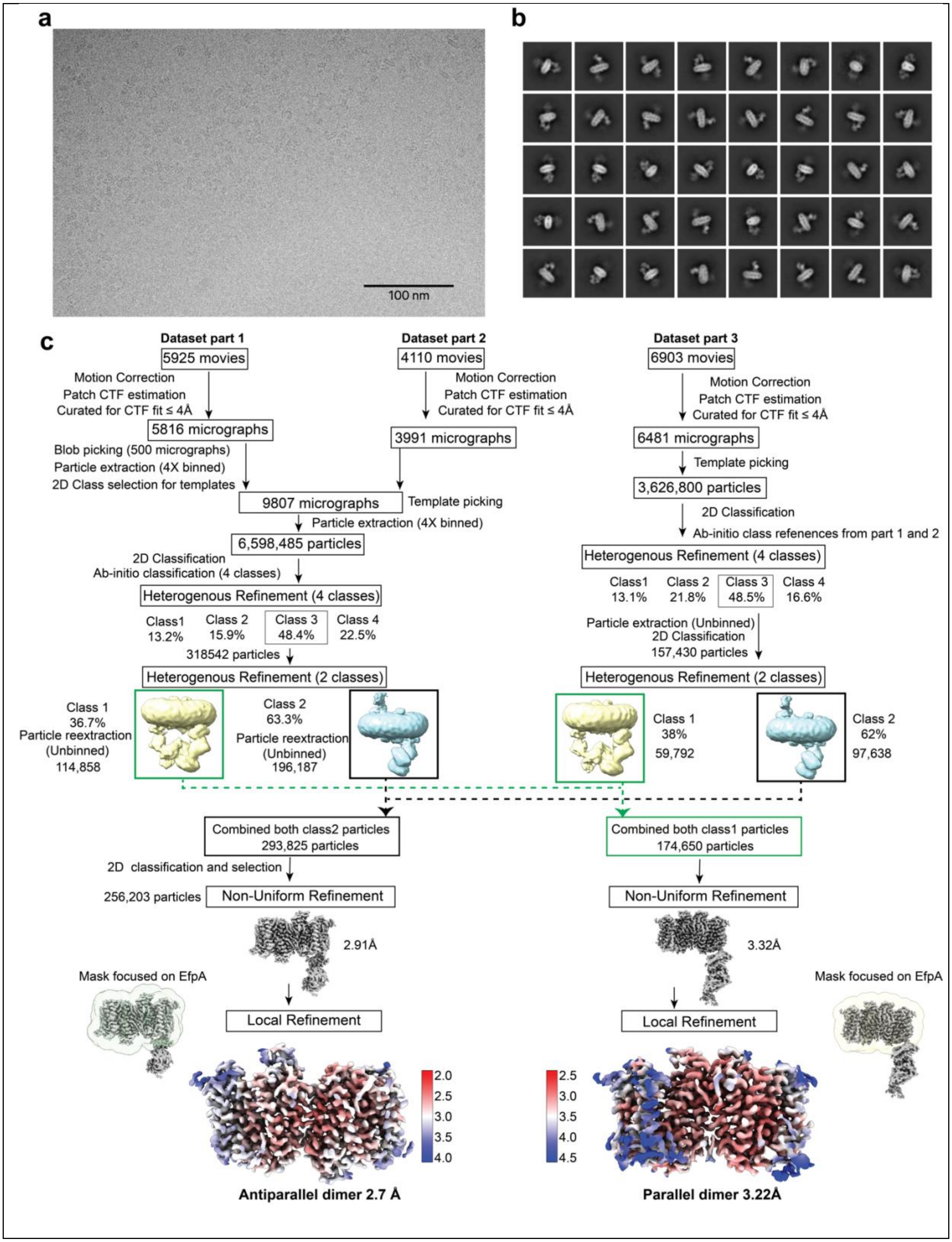
a
100 nm
b
c
Dataset part 1
5925 movies
Motion Correction
Patch CTF estimation
Curated for CTF fit ≤ 4Å
5816 micrographs
Blob picking (500 micrographs)
Particle extraction (4X binned)
2D Class selection for templates
Dataset part 2
4110 movies
Motion Correction
Patch CTF estimation
Curated for CTF fit ≤ 4Å
3991 micrographs
9807 micrographs
Template picking
Particle extraction (4X binned)
6,598,485 particles
2D Classification
Ab-initio classification (4 classes)
Heterogenous Refinement (4 classes)
Class1 13.2%
Class 2 15.9%
Class 3 48.4%
Class 4 22.5%
318542 particles
Heterogenous Refinement (2 classes)
Class 1 36.7% Particle reextraction (Unbinned) 114,858
Class 2 63.3% Particle reextraction (Unbinned) 196,187
Dataset part 3
6903 movies
Motion Correction
Patch CTF estimation
Curated for CTF fit ≤ 4Å
6481 micrographs
Template picking
3,626,800 particles
2D Classification
Ab-initio class refenences from part 1 and 2
Heterogenous Refinement (4 classes)
Class1 13.1%
Class 2 21.8%
Class 3 48.5%
Class 4 16.6%
Particle extraction (Unbinned)
2D Classification
157,430 particles
Heterogenous Refinement (2 classes)
Class 1 38% 59,792
Class 2 62% 97,638
Combined both class2 particles 293,825 particles
2D classification and selection
256,203 particles
Non-Uniform Refinement
2.91Å
Mask focused on EfpA
Local Refinement
2.0
2.5
3.0
3.5
4.0
Antiparallel dimer 2.7 Å
Combined both class1 particles 174,650 particles
Non-Uniform Refinement
3.32Å
Mask focused on EfpA
Local Refinement
2.5
3.0
3.5
4.0
4.5
Parallel dimer 3.22Å

**Extended Data Fig. 2| EfpA apo data processing workflow. a,** Representative micrograph image. **b,** Representative 2D classes form unbinned particles. **c,** Flowchart of 3D-map generation and refinement in cryoSPARC. Colored map representation of local resolution of the final map calculated from cryoSPARC. Left side shows the antiparallel dimer at global resolution 2.7 Å and right-side the parallel dimer at global resolution of 3.2 Å.

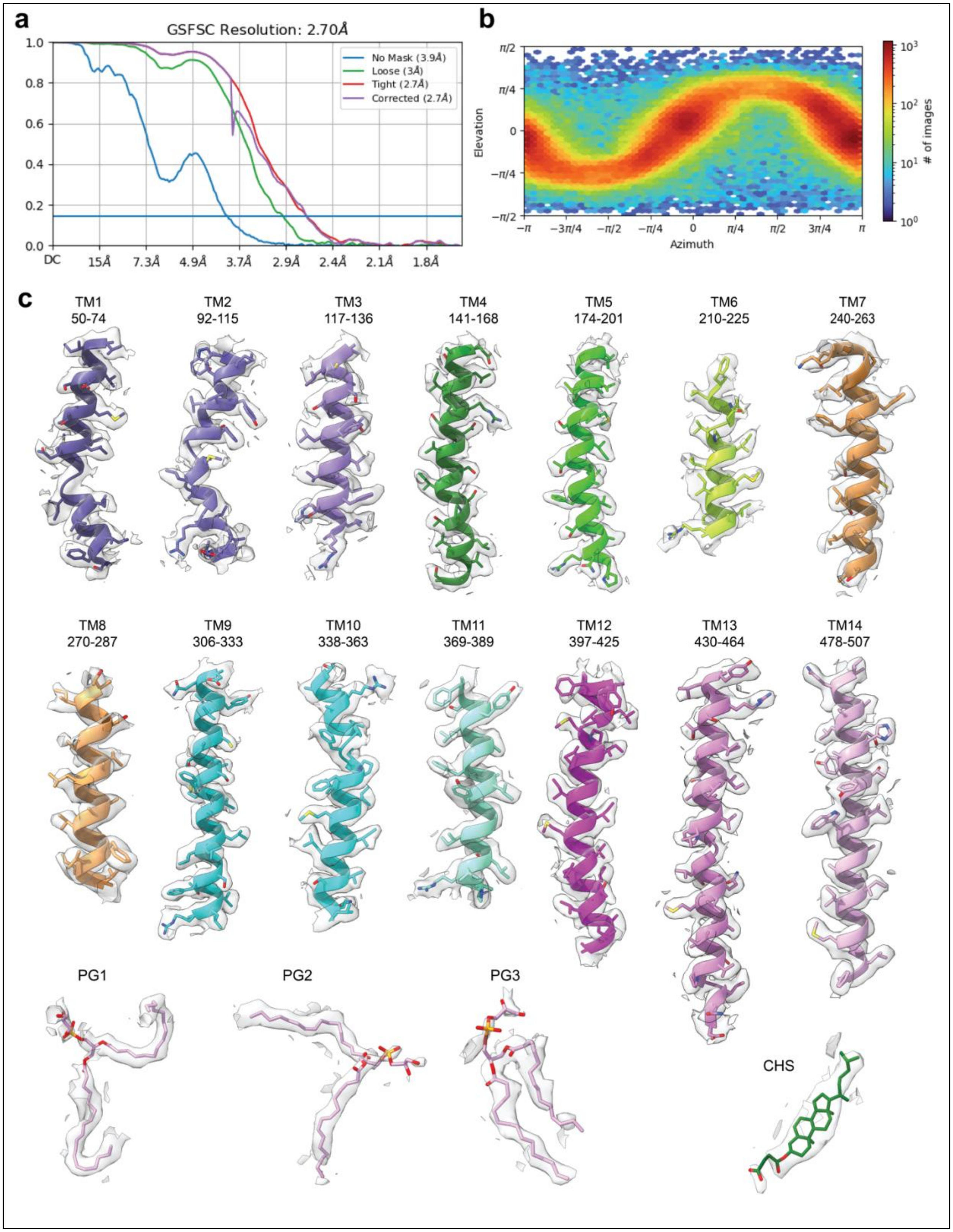
a
GSFSC Resolution: 2.70Å
No Mask (3.9Å)
Loose (3Å)
Tight (2.7Å)
Corrected (2.7Å)
DC
15Å
7.3Å
4.9Å
3.7Å
2.9Å
2.4Å
2.1Å
1.8Å
b
Elevation
Azimuth
### of images
c
TM1
50-74
TM2
92-115
TM3
117-136
TM4
141-168
TM5
174-201
TM6
210-225
TM7
240-263
TM8
270-287
TM9
306-333
TM10
338-363
TM11
369-389
TM12
397-425
TM13
430-464
TM14
478-507
PG1
PG2
PG3
CHS

**Extended Data Fig. 3| CryoEM density of antiparallel dimer apo map. a,** Fourier Shell Correlation (FSC) curve from final refinement showing a global resolution of 2.7Å at a threshold of 0.143. **b,** Euler angle heatmaps of EfpA apo antiparallel structure. **c,** Final model, and corresponding map densities for EfpA monomer transmembrane helices and associated three PG and one CHS molecules.

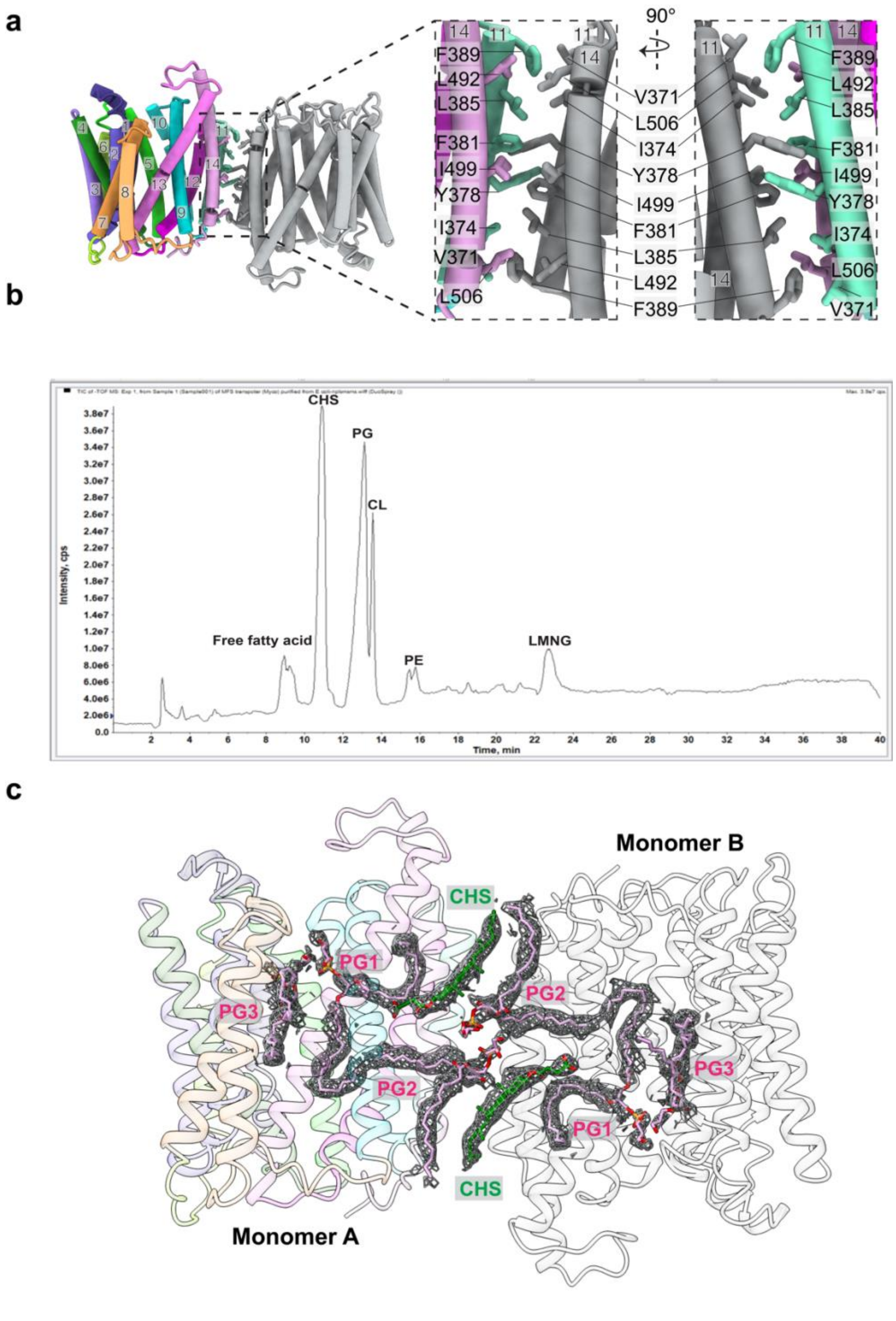


**Extended DATA Fig. 4| lipids in EfpA antiparallel dimer. a,** Residues involved in the interface between two monomers in the antiparallel dimer. **b,** EfpA lipid identification by mass spectrometry**. c,** cryoEM density for the lipids PG1, PG2, PG3 and CHS.

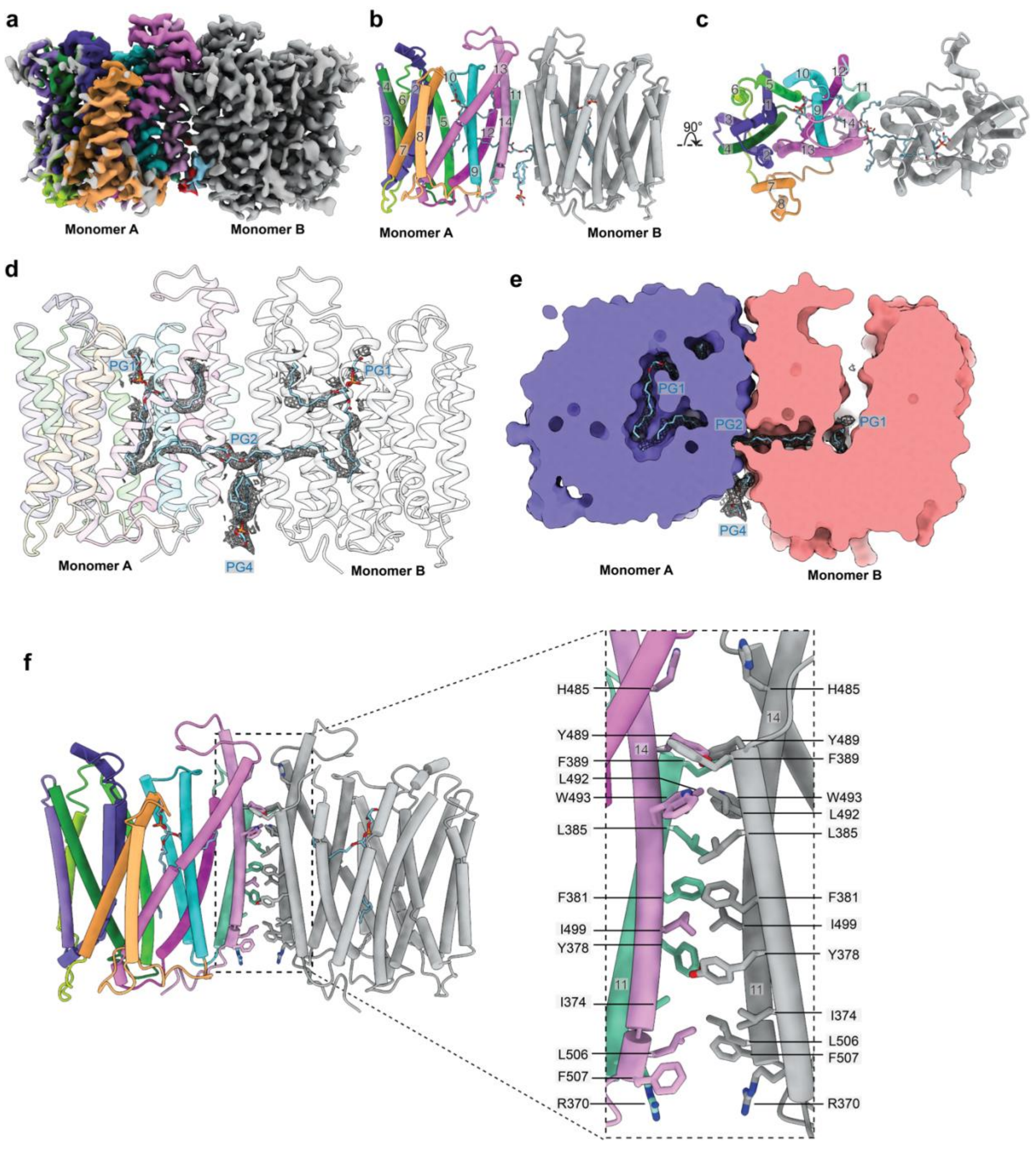


**Extended DATA Fig. 5|** Overall architecture of the parallel dimer **a,** Map for the parallel dimer. **b,** The structure of monomers A,B are schematized with lipids shown. **c,** top view as in b rotated by 90° from b, view down the two-fold axis between monomers. **d,** Density for the lipids PG1, PG2, PG4. **e,** Section through the solvent accessible surface showing regions occupied by lipids, and the outer vestibule. **f,** The interface between monomers showing residues involved in dimer formation.

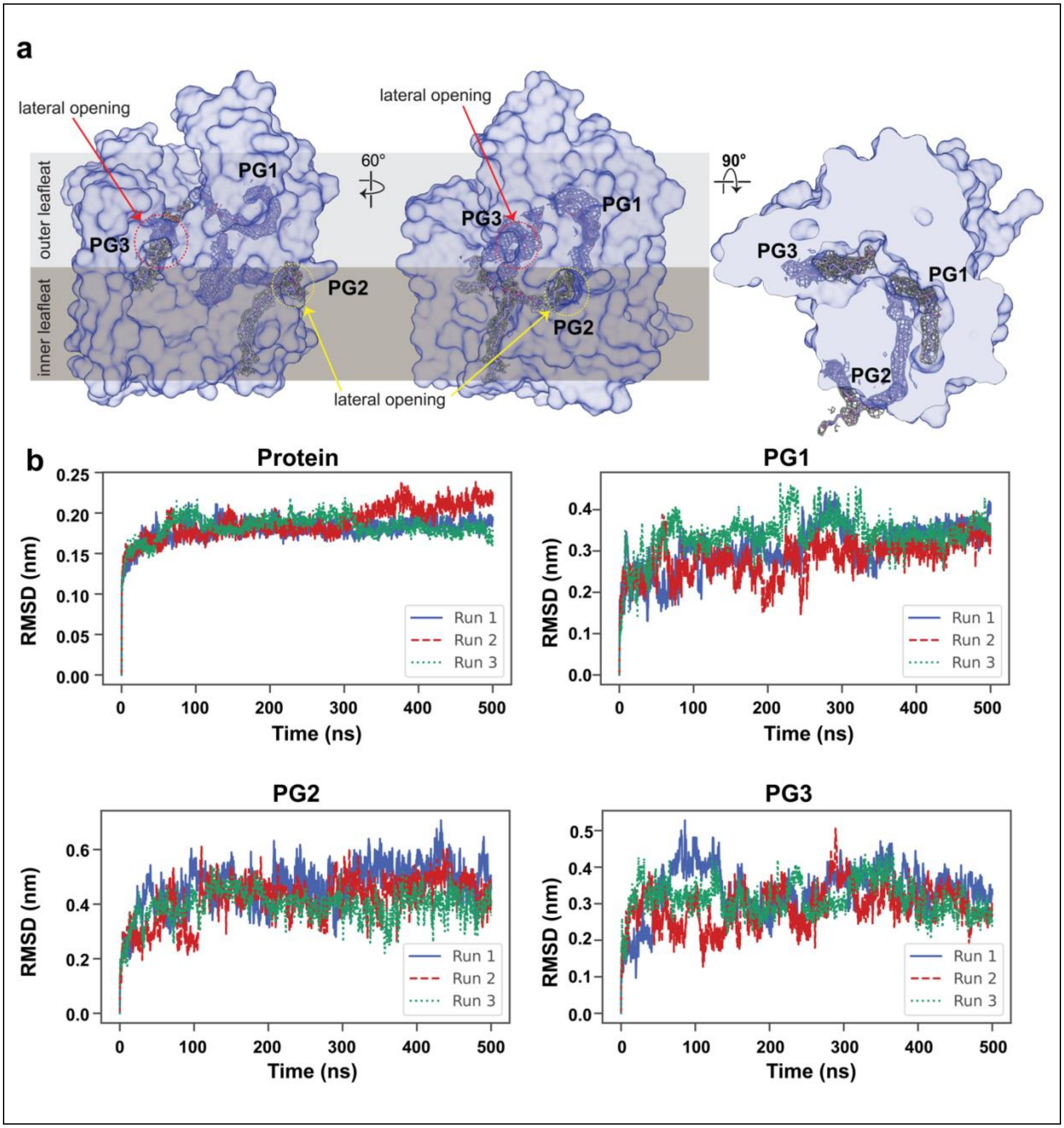

a
lateral opening
outer leafleat
inner leafleat
PG1
PG3
PG2
60°
90°
lateral opening
b
Protein
PG1
PG2
PG3
RMSD (nm)
Time (ns)
Run 1
Run 2
Run 3

**Extended DATA Fig. 6| Lipids molecules in EfpA monomer. a,** Lateral openings in EfpA apo monomer A. Left panel showing lateral opening towards the outer leaflet in red dotted circle with PG3 molecules and density of it in cryoEM map and lateral opening towards the inner leaflet in yellow dotted circle with PG2 molecules and density of it in cryoEM map. Middle panel is rotation of right panel at 80°. Right panel left panel rotated by 90° showing the PG1, PG2 and PG3 binding positions. **b,** MD simulations illustrating the binding characteristics and durability of the lipid molecules (PG1, PG2 and PG3) when interacting with the protein EfpA. The results (top left) reveal that the protein is stable with an RMSD of < 2.5 Å (displayed through the protein and Cα curves above) and a consistently stable binding of the lipid molecules is observed over a duration of 500 ns. This is evident from the fact that the PG1 and PG3 (top and bottom right) molecules exhibit an RMSD of < 5 Å while the PG2 molecule (bottom left) exhibits a more perturbed motion with a higher RMSD of < 7 Å.

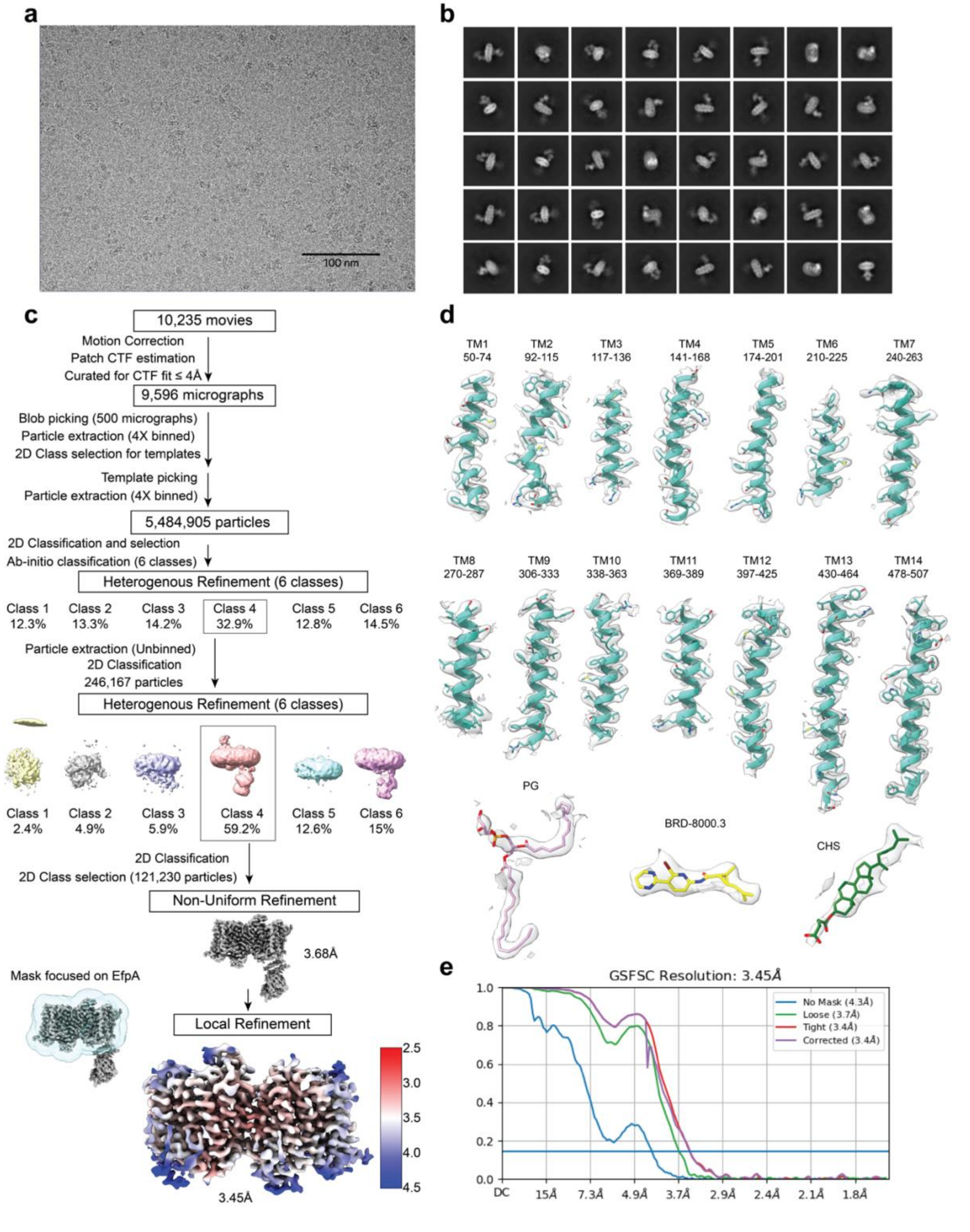

a
100 nm
b
c
10,235 movies
Motion Correction
Patch CTF estimation
Curated for CTF fit ≤ 4Å
9,596 micrographs
Blob picking (500 micrographs)
Particle extraction (4X binned)
2D Class selection for templates
Template picking
Particle extraction (4X binned)
5,484,905 particles
2D Classification and selection
Ab-initio classification (6 classes)
Heterogenous Refinement (6 classes)
Class 1 12.3%
Class 2 13.3%
Class 3 14.2%
Class 4 32.9%
Class 5 12.8%
Class 6 14.5%
Particle extraction (Unbinned)
2D Classification
246,167 particles
Heterogenous Refinement (6 classes)
Class 1 2.4%
Class 2 4.9%
Class 3 5.9%
Class 4 59.2%
Class 5 12.6%
Class 6 15%
2D Classification
2D Class selection (121,230 particles)
Non-Uniform Refinement
3.68Å
Mask focused on EfpA
Local Refinement
3.45Å
d
TM1 50-74
TM2 92-115
TM3 117-136
TM4 141-168
TM5 174-201
TM6 210-225
TM7 240-263
TM8 270-287
TM9 306-333
TM10 338-363
TM11 369-389
TM12 397-425
TM13 430-464
TM14 478-507
PG
BRD-8000.3
CHS
e
GSFSC Resolution: 3.45Å
No Mask (4.3Å)
Loose (3.7Å)
Tight (3.4Å)
Corrected (3.4Å)
DC 15Å 7.3Å 4.9Å 3.7Å 2.9Å 2.4Å 2.1Å 1.8Å

**Extended DATA Fig. 7| Workflow of EfpA BRD-8000.3 bound data processing. a,** Representative micrograph image. **b,** Representative 2D classes form unbinned particles extract. **c,** Flowchart of 3D-map generation and refinement in cryoSPARC. Colored map for representation of local resolution of the final map calculated from cryoSPARC. **d,** Model and corresponding densities for EfpA monomer transmembrane helices and BRD-8000.3 and PG lipid molecule. **e,** Fourier Shell Correlation (FSC) curve from final refinement showing a global resolution of 3.45Å at a threshold of 0.143.

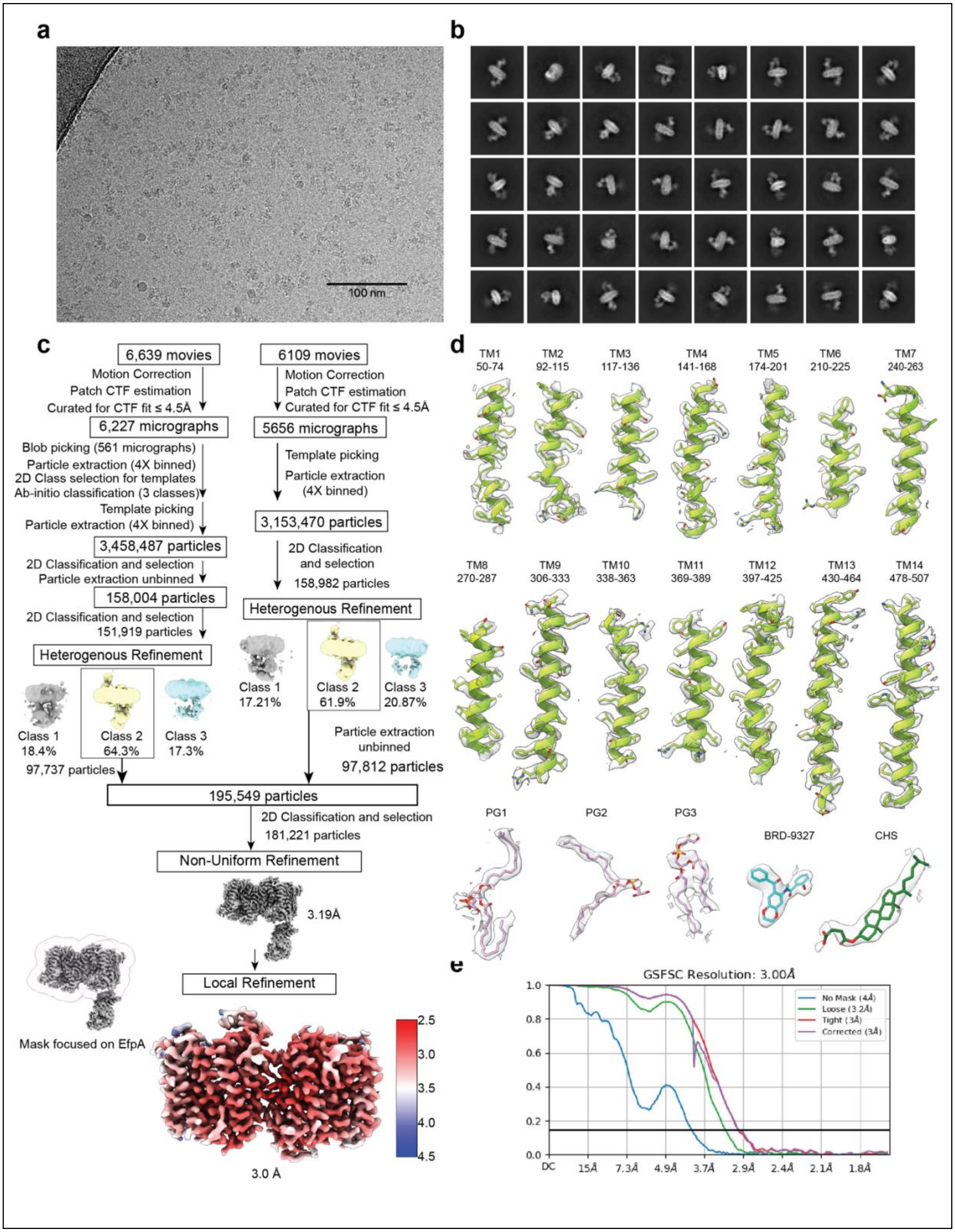

a
100 nm
b
c
6,639 movies
Motion Correction
Patch CTF estimation
Curated for CTF fit ≤ 4.5Å
6,227 micrographs
Blob picking (561 micrographs)
Particle extraction (4X binned)
2D Class selection for templates
Ab-initio classification (3 classes)
Template picking
Particle extraction (4X binned)
3,458,487 particles
2D Classification and selection
Particle extraction unbinned
158,004 particles
2D Classification and selection
151,919 particles
Heterogenous Refinement
Class 1 18.4%
Class 2 64.3%
Class 3 17.3%
97,737 particles
6109 movies
Motion Correction
Patch CTF estimation
Curated for CTF fit ≤ 4.5Å
5656 micrographs
Template picking
Particle extraction (4X binned)
3,153,470 particles
2D Classification and selection
158,982 particles
Heterogenous Refinement
Class 1 17.21%
Class 2 61.9%
Class 3 20.87%
Particle extraction unbinned
97,812 particles
195,549 particles
2D Classification and selection
181,221 particles
Non-Uniform Refinement
3.19Å
Local Refinement
Mask focused on EfpA
2.5
3.0
3.5
4.0
4.5
3.0 Å
d
TM1 50-74
TM2 92-115
TM3 117-136
TM4 141-168
TM5 174-201
TM6 210-225
TM7 240-263
TM8 270-287
TM9 306-333
TM10 338-363
TM11 369-389
TM12 397-425
TM13 430-464
TM14 478-507
PG1
PG2
PG3
BRD-9327
CHS
e
GSFSC Resolution: 3.00Å
No Mask (4Å)
Loose (3.2Å)
Tight (3Å)
Corrected (3Å)
1.0
0.8
0.6
0.4
0.2
0.0
DC
15Å
7.3Å
4.9Å
3.7Å
2.9Å
2.4Å
2.1Å
1.8Å

**Extended DATA Fig. 8| Workflow of EfpA BRD-9327 bound cryoEM data processing. a,** Representative micrograph image. **b,** Representative 2D classes form unbinned particles extract. **c,** Flowchart of 3D-map generation and refinement in cryoSPARC . Colored map for representation of local resolution of the final map calculated from cryoSPARC. **d,** Model and corresponding densities for EfpA monomer transmembrane helices and BRD-8000.3 and PG lipid molecule. **e,** Fourier Shell Correlation (FSC) curve from final refinement showing a global resolution of 3.0Å at a threshold of 0.143.

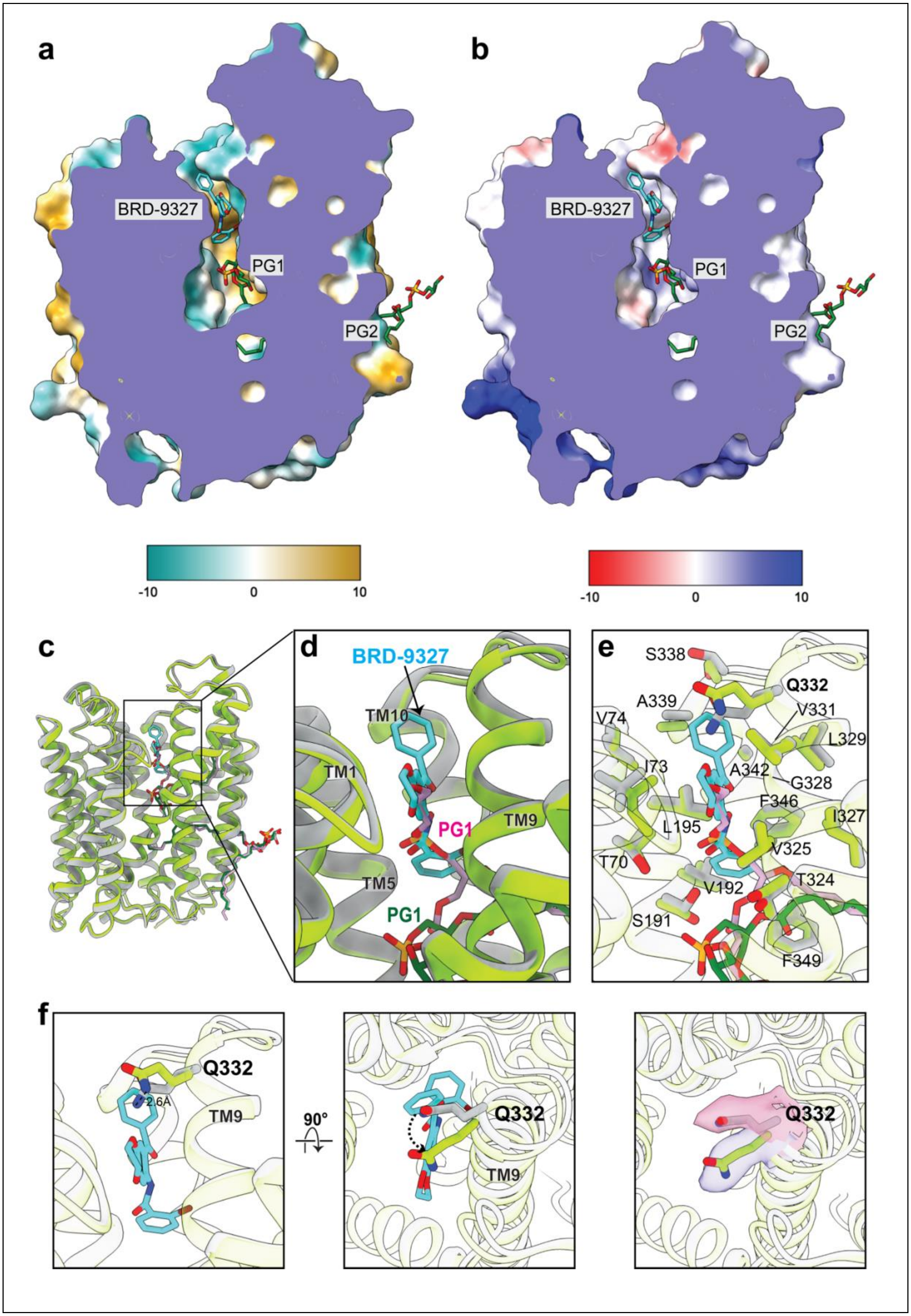
a
b
BRD-9327
PG1
PG2
BRD-9327
PG1
PG2
-10
0
10
-10
0
10
c
d
BRD-9327
TM10
TM1
PG1
TM9
TM5
PG1
e
S338
Q332
A339
V331
V74
L329
I73
A342
G328
F346
I327
L195
V325
T70
T324
V192
S191
F349
f
Q332
2.6A
TM9
90°
Q332
TM9
Q332

**Extended DATA Fig. 9| Structural comparison of EfpA apo and BRD-9327 bound structures. a,** section of the electrostatic surface of BRD-9327-bound EfpA showing the BRD-9327 binding site. **b**, as in a, colored according to hydrophobicity. **c,** superposition of BRD-9327 (green) onto the apo structure (gray) with lipids PG1 and PG2 (pink) as in the apo structure. BRD-9327 bound structure PG1 and PG2 shown in dark green and BRD-9327 (cyan) with nitrogen (blue), oxygen (red), and phosphate (orange). **d,** the BRD-9327 binding site and comparison with the apo structure, showing the overlap between PG1 of apo and BRD-9327. **e,** side chains of binding site shown in d. **f,** outward movement of the Q332 side chain in the BRD-9327 bound structure. Middle panel after rotating the structure 90° showing the top view. Right panel shows the EM density around the Q332 for BRD-9327 bound structure (blue) and apo structure (pink).

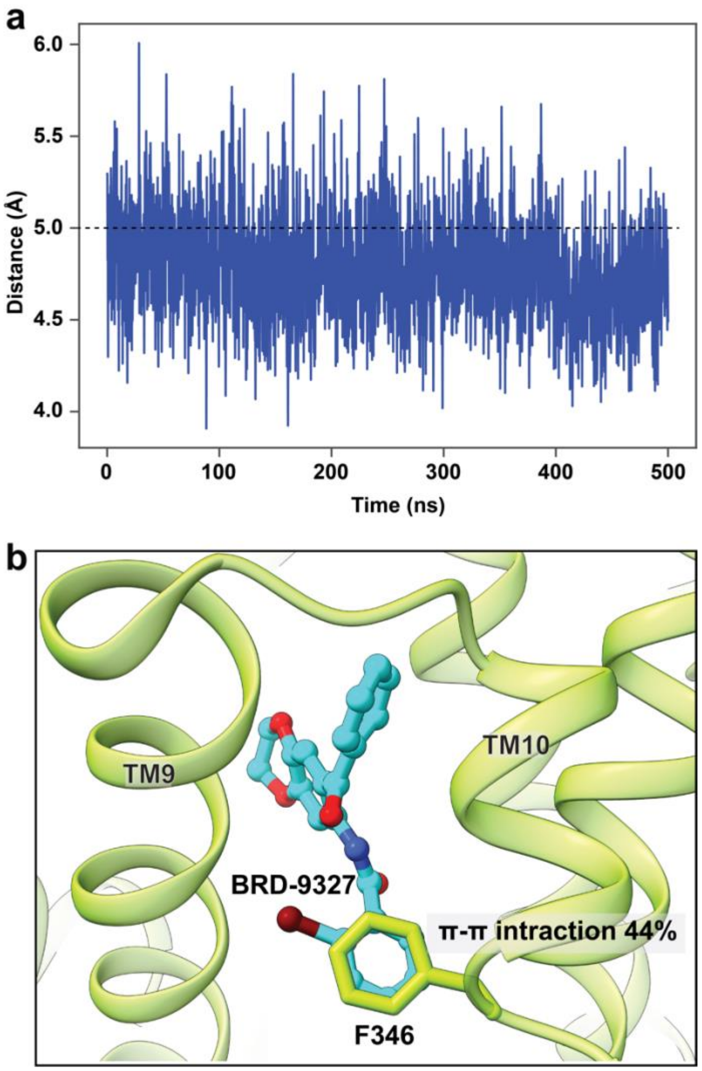


**Extended Data Fig. 10| Molecular dynamics simulations of BRD-9327 interaction with EfpA. a,** distances between center of mas of BRD-9327 benzyl bromide ring and F346 side chain ring in 500 ns MD simulations. Distances less than 5 Å indicate π –π interactions (black dashed line). **b,** Representative snapshots (187 ns) for MD simulations of BRD-9327 binding to EfpA showing π–π interactions between BRD-9327 and $F346^{TM10}$. The π–π interactions between BRD-9327 and $F346^{TM10}$ were maintained for more ~ 44% throughout the simulation.

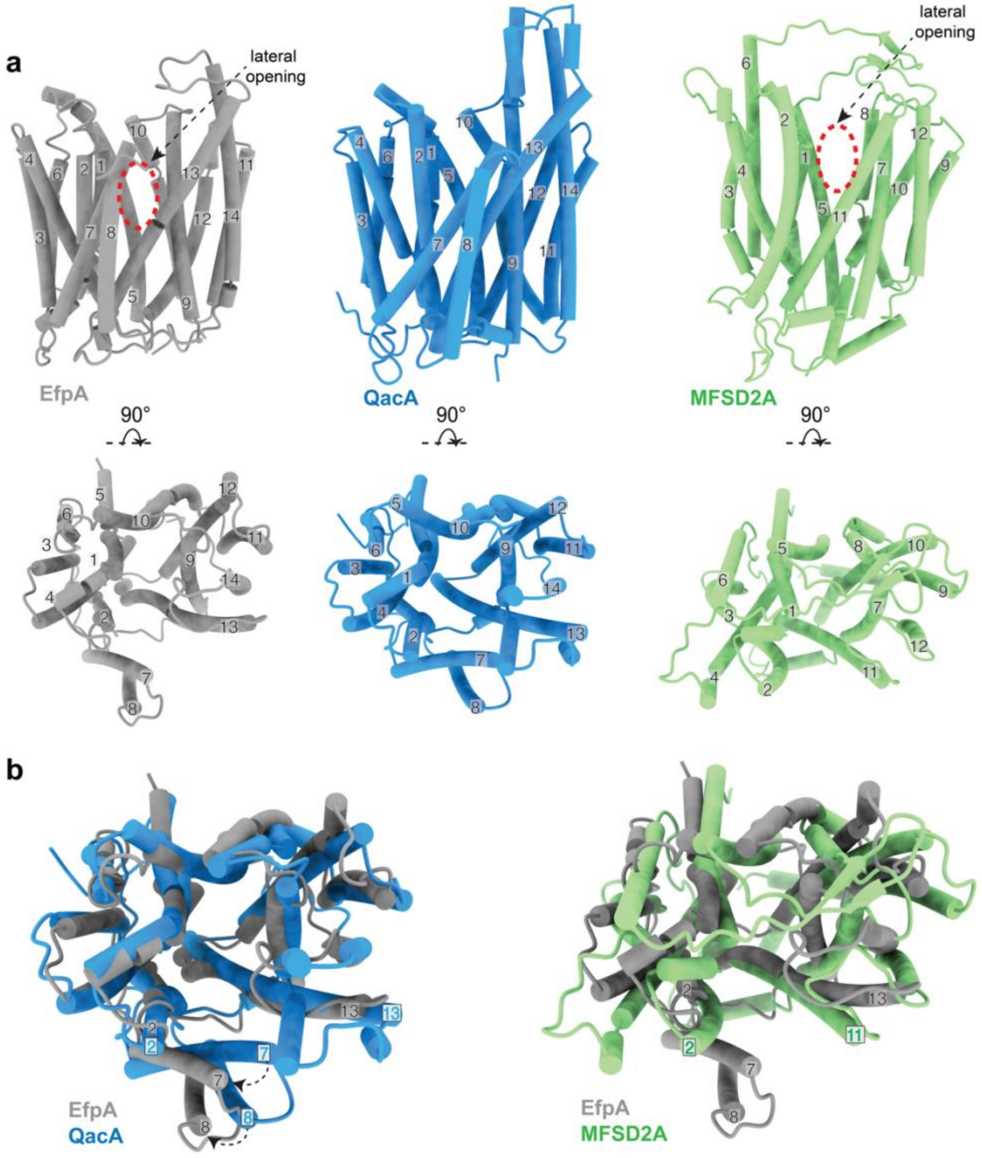


**Extended Data Fig. 11| Comparison of EfpA architecture with other DHA2 family transporters (QacA and NorC) and lipid transporter MFSD2A. a,** Top panel, side view and bottom panel rotated by 90° to show the top view of the TM arrangement in EfpA, QacA (PDB: 7Y58) and MFSD2A (PDB:7N98), The extra linker between TM7 and TM8 in quaternary ammonium compound A family members (EfpA and QacA) lies next to the lateral opening between TM2 and TM13. EfpA TM7 and 8 arrange opposite towards TM2 in way that they don't block this lateral opening. In QacA TM7 and 8 present juxtaposed to space between TM2 and 13 and block the lateral opening. The absence of an extra linker domain in MFSD2A results in lateral opening similar to EfpA. **b,** Top view of overlapping structures of EfpA with QacA and MFSD2A highlighting the rearrangement of TM7 and 8.

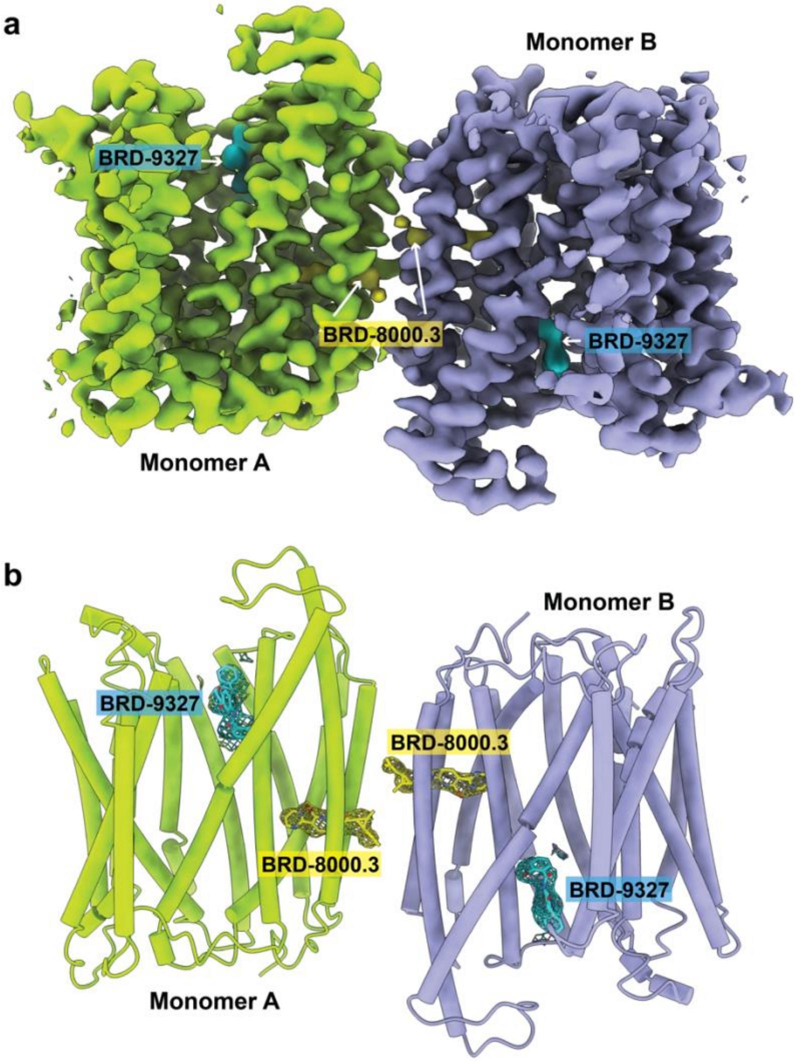


**Extended Data Fig. 12| Combinatorial inhibition of EfpA. a,** cryo-EM map density of BRD-9327 and BRD-8000.3 bound EfpA$^{EM}$. Both monomers of EfpA shown as yellow-green and violet color respectively. Each monomer has one BRD-9327 molecule (aqua color) and one BRD-8000.3 (yellow color). **b,** Tertiary structure of EfpA$^{EM}$ monomer A with BRD-9327 (aqua) and BRD-8000.3 (yellow color) bound. Density for BRD-9327 (aqua) and BRD-8000.3 (yellow color) are shown as mesh. Color by atom Nitrogen (blue), oxygen (red) and bromide (brown).

| Extended Data Table 1 | | | | |
|---|---|---|---|---|
| **Structure** | EfpA (antiparallel dimer) | EfpA (parallel dimer) | BRD-8000.3 bound EfpA | BRD-9327 bound EfpA |
| **PDB** | **9BII** | **9BL7** | **9BIN** | **9BIQ** |
| **EMDB** | **44591** | **44651** | **44594** | **44598** |
| **Data collection / processing** | | | | |
| Magnification | 105,000 | 105,000 | 105,000 | 105,000 |
| Voltage (kV) | 300 | 300 | 300 | 300 |
| Pixel size (Å) | 0.835 | 0.835 | 0.4175 | 0.417 |
| Pixel size after binning (Å) | 0.835 | 0.835 | 0.835 | 0.834 |
| Defocus range (µm) | -1 to -2 | -1 to -2 | -1 to -2 | -1 to -2 |
| Electron exposure (e/Å$^2$) | ~45 | ~45 | ~45 | ~43 |
| Symmetry imposed | C1 | C1 | C1 | C1 |
| Initial particles (No.) | 10,225,285 | 10,225,285 | 5,484,905 | 6,611,957 |
| Final particles (No.) | 256,203 | 174,650 | 121,230 | 181,221 |
| Map resolution (Å) | 2.7 | 3.22 | 3.45 | 3.0 |
| FSC threshold | 0.143 | 0.143 | 0.143 | 0.143 |
| Map resolution range (Å) | 2.3 – 33.1 | 2.7 – 39.3 | 2.9 – 12.6 | 2.5 – 29.3 |
| **Refinement** | | | | |
| Initial model used | AF-P9WJY4-F1 | | | |
| Model resolution (Å) | 2.9 | 3.39 | 3.6 | 3.2 |
| FSC threshold | 0.5 | 0.5 | 0.5 | 0.5 |
| Map sharpening B factor (Å$^2$ ) | 88.2 | 91.7 | 119.2 | 101 |
| Model composition | | | | |
| Non-hydrogen atoms | 7224 | 7059 | 7086 | 7283 |
| Protein residues | 940 | 940 | 942 | 942 |
| Ligands | - | - | 2 | 2 |
| Lipids | 6 | 4 | 2 | 6 |

| | | | | |
|---|---|---|---|---|
| B factors (Å$^2$) | | | | |
| Protein | 48.61 | 53.68 | 76.39 | 45.29 |
| Ligand/Lipids | 37.39 | 46.54 | 37.87 | 20 |
| RMS deviations | | | | |
| Bond lengths (Å) | 0.006 | 0.003 | 0.005 | 0.008 |
| Bond angles (°) | .607 | 0.535 | 0.558 | 0.690 |
| Validation | | | | |
| MolProbity score | 1.20 | 1.49 | 1.53 | 1.33 |
| Clashscore | 2.68 | 4.76 | 5.46 | 4.14 |
| Poor rotamers (%) | 0 | 0 | 0 | 0 |
| Ramachandran plot (%) | | | | |
| Favored (%) | 2.78 | 3.63 | 3.53 | 2.67 |
| Allowed (%) | 97.22 | 96.37 | 96.47 | 97.33 |
| Disallowed (%) | 0 | 0 | 0 | 0 |
